# Super-resolved protein imaging using bifunctional light-up aptamers

**DOI:** 10.1101/2024.01.26.577321

**Authors:** Franziska Grün, Niklas van den Bergh, Maja Klevanski, Mrigank S. Verma, Bastian Bühler, G. Ulrich Nienhaus, Thomas Kuner, Andres Jäschke, Murat Sunbul

## Abstract

Efficient labeling methods for protein visualization with minimal tag size and appropriate photophysical properties are required for single-molecule localization microscopy (SMLM), providing insights into the organization and interactions of biomolecules in cells at the molecular level. Among the fluorescent light-up aptamers (FLAPs) originally developed for RNA imaging, RhoBAST stands out due to its remarkable brightness, photostability, fluorogenicity, and rapid exchange kinetics, enabling super-resolved imaging with high localization precision. Here, we expand the applicability of RhoBAST to protein imaging by fusing it to protein-binding aptamers. The versatility of such bifunctional aptamers is demonstrated by employing a variety of protein-binding DNA or RNA aptamers and different FLAPs. Moreover, fusing RhoBAST with the GFP-binding aptamer AP3 facilitates high- and super-resolution imaging of GFP-tagged proteins, which is particularly valuable in view of the widespread availability of plasmids and stable cell lines expressing proteins fused to GFP. The bifunctional aptamers compare favorably with standard antibody-based immunofluorescence protocols, as they are 7-fold smaller than antibody conjugates and exhibit higher bleaching-resistance. We demonstrate the effectiveness of our approach in super-resolution microscopy in secondary mammalian cell lines and primary neurons by RhoBAST-PAINT, an SMLM protein imaging technique that leverages the transient binding of the fluorogenic rhodamine dye SpyRho to RhoBAST.

## Introduction

In single-molecule localization microscopy (SMLM), the activation, photoswitching or transient binding of individual fluorophores coupled with the determination of their positions during the fluorescent event enables super-resolved fluorescence imaging beyond the diffraction limit.^[1]^ While conventional fluorescence microscopy has provided insights into the organization of cellular compartments and tissue composition, SMLM has advanced our understanding of cellular functions and biomolecular interactions to the molecular level.^[2]^ SMLM encompasses various techniques, including PALM (photo-activated localization microscopy),^[3]^ *d*STORM (direct stochastic optical reconstruction microscopy),^[4]^ as well as DNA-PAINT (DNA point accumulation for imaging in nanoscale topography)^[5]^ to investigate protein nanoarchitectures with nanometer-scale resolution.

In PALM, photoactivatable fluorescent proteins are genetically fused to the protein of interest (POI) and expressed in cells.^[3]^ In contrast, small organic fluorophores have been employed for protein visualization through the use of the genetically encodable HaloTag and SNAP-tag.^[6],[7]^ These self-labeling and genetically encodable protein tags covalently attach the fluorophores to the POI within the cell.^[6],[7]^

If genetic modification shall be avoided, immunostaining remains the most common technique for visualizing proteins in fixed cells and tissues. Typically, a primary antibody binds to the POI and is visualized with a fluorophore-labeled secondary antibody, binding the primary antibody. Immunostaining for super-resolution microscopy utilizes antibodies^[4]^ or nanobodies^[8]^ linked to photoswitchable fluorophores in *d*STORM. Alternatively, this advancement has also been achieved by employing antibodies labeled with a DNA “docking strand” that can hybridize to fluorophore-labeled DNA “imager strand” in DNA-PAINT.^[9]^ However, conjugates of primary and secondary antibodies constitute relatively large tags compared to RNA or DNA aptamers. Antibodies directly labeled with fluorophores are also used, but the fluorophore conjugation can affect their binding to the POI.^[10]^ Single-stranded oligonucleotide aptamers exhibit selective and high-affinity binding to their targets, including small molecules^[11]^ or proteins.^[12]^ Protein-binding aptamers were utilized in DNA-PAINT by conjugation of the aptamer to an oligonucleotide docking strand. Examples include slow off-rate modified aptamers (SOMAmers) targeting various proteins^[13]^ or the GFP-binding AP3 aptamer.^[14]^ Further RNA aptamers were selected to specifically bind fluorogenic dyes with high affinity paired with fluorescence light-up.^[11, 15-19]^ These genetically encodable tags, known as fluorescent light-up aptamers (FLAPs), find application in visualizing and tracking of RNAs in live cells.^[17, 20, 21]^ Compared to other FLAPs, RhoBAST is particularly suitable for SMLM due to its fast dye exchange kinetics, high brightness, and outstanding bleaching resistance.^[19]^ Its ligand was further optimized, yielding the fluorogenic spirocyclic rhodamine SpyRho.^[22]^

This work introduces a novel protein labeling method based on bifunctional aptamers for super-resolution imaging. Specifically, the approach involves fusing different protein-binding aptamers with RhoBAST. The resulting bifunctional aptamers bind to the respective POIs, whereupon the intermittent fluorescence, crucial for SMLM, is achieved through the interaction of SpyRho with RhoBAST (Fig. 1A). We implemented this design with various protein-binding aptamers, including a GFP-binding aptamer for SMLM imaging of GFP-tagged proteins, as well as diverse aptamers binding to endogenous target proteins. Furthermore, live-cell imaging of GFP-tagged proteins was achieved with genetically encoded AP3-RhoBAST and cell-permeable SpyRho. The bifunctional aptamers were employed in super-resolved protein imaging, and we compared their performance with commercially available antibodies.

**Figure 1:**
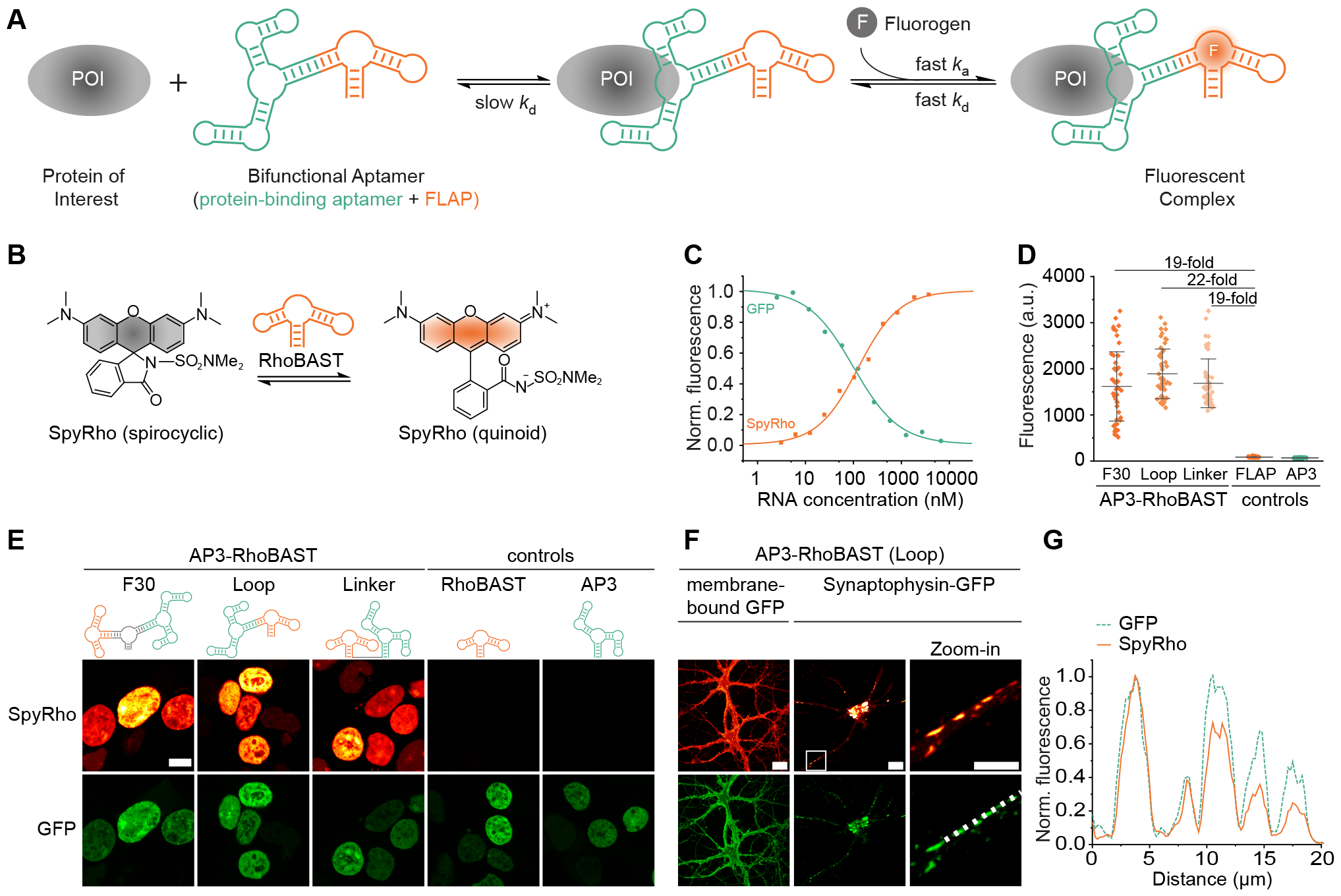
Characterization of bifunctional aptamer AP3-RhoBAST for protein visualization in fixed cells. **A)** Schematic drawing of a bifunctional aptamer consisting of a protein-binding aptamer (green) and a fluorescent light-up aptamer (FLAP, orange). The protein-binding aptamer binds the POI, while the FLAP binds a fluorogen, resulting in intermittent fluorescence due to fast association (*k*_a_) and dissociation (*k*_d_) rate coefficients. **B)** Equilibrium of spirocyclic rhodamine (SpyRho) between the colorless, non-fluorescent spirocyclic form and the colored, fluorescent quinoid form. SpyRho lights up upon binding to the aptamer, RhoBAST. **C)** Binding isotherms of AP3-RhoBAST (loop construct) with SpyRho (10 nM, orange line) and GFP (5 nM, green line) giving dissociation constants of *K*_D_ = 107 ± 17 nM and *K*_D_ = 61 ± 8 nM, respectively. Minimal and maximal fluorescence intensities were normalized to 0 and 1, respectively. *K*_D_-values are given as mean ± s.d., *N* = 3 independent experiments. **D)** Quantification of SpyRho fluorescence intensities (mean ± s.d., *N* = 50 cells) of HEK293T cells expressing H2B-GFP, derived from confocal images as shown in E) giving the *in cellulo* fluorescence turn-on values. **E)** Confocal images of fixed and permeabilized HEK293T cells expressing H2B-GFP. Cells were incubated with 500 nM aptamer, washed and stained with 100 nM SpyRho. Scale bar, 20 µm and 10 µm for zoom-in. **F)** Confocal images of fixed and permeabilized neurons expressing membrane-bound GFP-tag or synaptophysin-GFP. Cells were incubated with 500 nM AP3-RhoBAST, washed and incubated with 100 nM SpyRho prior to imaging. White frame indicates the region of interest, depicted as zoom-in. Scale bars, 20 µm. **G)** Normalized fluorescence intensity profiles of GFP (green, dashed line) and SpyRho (orange, solid line) along the dashed line in F).

## Results and Discussion

### Exploring bifunctional RhoBAST aptamers for protein visualization

The RhoBAST:SpyRho system offers remarkable features such as high quantum yield (QY = 0.95), excellent photostability, and rapid association and dissociation kinetics for super-resolved imaging of biomolecules using SMLM.^[22]^ By binding to RhoBAST, SpyRho gains an intrinsic resistance to photobleaching that surpasses even the state-of-the-art fluorophore JF549. This feature, combined with the high brightness, allows for collecting a large number of photons per blinking event, thus increases the localization precision. The fast and continuous ligand exchange of RhoBAST:SpyRho enables rapid acquisition of super-resolved images with localization precision not limited by photobleaching. Super-resolved imaging facilitated by RhoBAST:SpyRho using SMLM adheres to principles analogous to DNA-PAINT. However, the fluorogenic property of SpyRho -a fluorescence turn-on of about 60-fold upon binding to RhoBAST - is a definitive advantage of RhoBAST:SpyRho over traditional DNA-PAINT, providing a higher signal to background ratio.^[22]^

To enable straightforward and widely applicable protein imaging by SMLM, we sought to leverage the capabilities of RhoBAST to protein-binding aptamers by creating bifunctional aptamers (Fig. 1A). For proof-of-principle experiments, we first selected GFP-tagged proteins as target proteins and AP3 as the protein-binding RNA aptamer, which specifically binds members of the GFP-family including EGFP, CFP, and YFP with high affinities.^[23]^ Incubation of cells expressing GFP-tagged target proteins with the bifunctional AP3-RhoBAST fusion aptamer would enable labeling of the target protein with RhoBAST. Colocalization of GFP and RhoBAST fluorescent signals in the presence of SpyRho serves as a valuable tool for verifying the functionality and specificity of AP3-RhoBAST. Furthermore, considering the vast number of expression plasmids coding for GFP-tagged proteins and cell lines stably expressing GFP-tagged proteins, it is attractive to devise a method that leverages the GFP tag for enabling super-resolution imaging by SMLM.

To create the bifunctional aptamer AP3-RhoBAST, we tested three different scaffolds for linking AP3 and RhoBAST: i) The F30-scaffold (F30),^[24]^ characterized by a stable three-way junction, supporting stable and independent folding of both connected aptamers; ii) insertion of the protein-binding aptamer into RhoBAST’s tetraloop (loop), known to be exchangeable without compromising the binding affinity or specificity to SpyRho;^[22]^ iii) a flexible, single-stranded 10-nucleotide linker (linker) between the aptamers. All bifunctional and individual aptamers were *in vitro* transcribed, purified by polyacrylamide gel electrophoresis, and *in vitro* characterized. The binding affinities between the bifunctional aptamers and SpyRho (or GFP) were determined by measuring the fluorescence change while titrating the aptamers to SpyRho (or GFP). SpyRho lights up upon interaction with RhoBAST, whereas AP3 binding decreases GFP’s fluorescence (Fig. 1C). Notably, SpyRho showed high binding affinities towards all bifunctional AP3-RhoBAST scaffolds, with *K*D values of 122 nM (F30), 107 nM (Loop), and 119 nM (Linker) (Fig. 1C, Fig. S1B, C, Table S1). Furthermore, all designed bifunctional AP3-RhoBAST constructs exhibited a high binding affinity to GFP, with *K*D values ranging from 61 to 89 nM (Fig. 1C, Fig. S1B, C). Moreover, fusing RhoBAST with AP3 did not substantially affect the light-up of the RhoBAST:SpyRho complex, shown by high fluorescence turn-on values with all tested constructs (Table S1).

Next, we investigated the *in cellulo* performance of the bifunctional aptamers using confocal laser-scanning microscopy (CLSM). HEK293T cells expressing GFP-tagged histone H2B protein were fixed, permeabilized, incubated with the freshly folded AP3-RhoBAST aptamer, and then washed. Prior to imaging, cells were incubated with SpyRho. In confocal images, we exclusively detected the SpyRho fluorescence within the nucleus, consistent with the nuclear localization of H2B. Colocalization of GFP and SpyRho fluorescence verified the specificity of the AP3-RhoBAST and GFP-H2B interaction (Fig. 1E, Fig. S1D-F). In contrast, the individually applied RhoBAST or AP3 aptamers, employed as negative controls, yielded no fluorescent signal (Figure 1D, E). Across all scaffold designs of AP3-RhoBAST, H2B-GFP visualization *in cellulo* was achieved with comparable turn-on ratios. The quantification of the SpyRho fluorescent signal revealed substantial signal-to-background ratios of 19 to 22 (Fig. 1D). Due to its best signal-to-background ratio, most compact predicted secondary structure and shortest sequence, AP3-RhoBAST fused via the tetraloop of RhoBAST was selected for all subsequent experiments (Fig. S1G).

Next, we expanded the application of AP3-RhoBAST to primary cells expressing GFP-tagged proteins. To this end, rat hippocampal neurons were transduced, using adeno-associated virus, to transiently express the major synaptic vesicle protein, synaptophysin-GFP, and a membrane-bound GFP-tag. These neurons were labeled with AP3-RhoBAST and imaged in the presence of SpyRho as described before. Visualization by confocal microscopy and the observed colocalization of GFP and SpyRho fluorescence revealed the successful targeting of synaptophysin-GFP in neurites and cell soma as well as membrane-bound GFP at the plasma membrane of neurons by AP3-RhoBAST and SpyRho (Fig. 1F, G, Fig. S2).

### Bifunctional RhoBAST aptamers for imaging endogenous proteins in fixed and on live cells

To visualize endogenous proteins within cells, RhoBAST can be combined with a protein-binding aptamer directly targeting the protein of interest. In these experiments, we utilized the single-stranded DNA aptamer AS1411, which targets nucleolin, a highly abundant protein found in the nucleolus.^[25, 26]^ Due to its elevated expression in several cancer types, nucleolin serves as an attractive target for both diagnostic and therapeutic purposes, with AS1411 previously explored in drug-conjugate studies and as a therapeutic agent.^[27]^ Here, we exploited AS1411 to visualize endogenous nucleolin in cells. To create bifunctional AS1411-RhoBAST as a DNA-RNA conjugate, we ligated the 3’ end of *in vitro* transcribed RhoBAST with the 5’ end of single-stranded AS1411 oligonucleotide (Fig. S3). The resulting bifunctional AS1411-RhoBAST aptamer was purified by polyacrylamide gel electrophoresis and characterized. As expected, AS1411-RhoBAST retained RhoBAST’s high affinity to SpyRho (*K*D = 113 ± 4 nM) and a large fluorescence turn-on ratio of 37-fold. Confocal imaging of endogenous nucleolin in fixed and permeabilized HEK293T cells with AS1411-RhoBAST and SpyRho showed successful labeling, as indicated by the localized SpyRho fluorescence in the nucleolus (Fig. 2A). In control samples, where either the RhoBAST or AS1411 aptamers were used alone instead of the bifunctional aptamer, we did not observe any specific fluorescent pattern (Fig. 2A).

**Figure 2:**
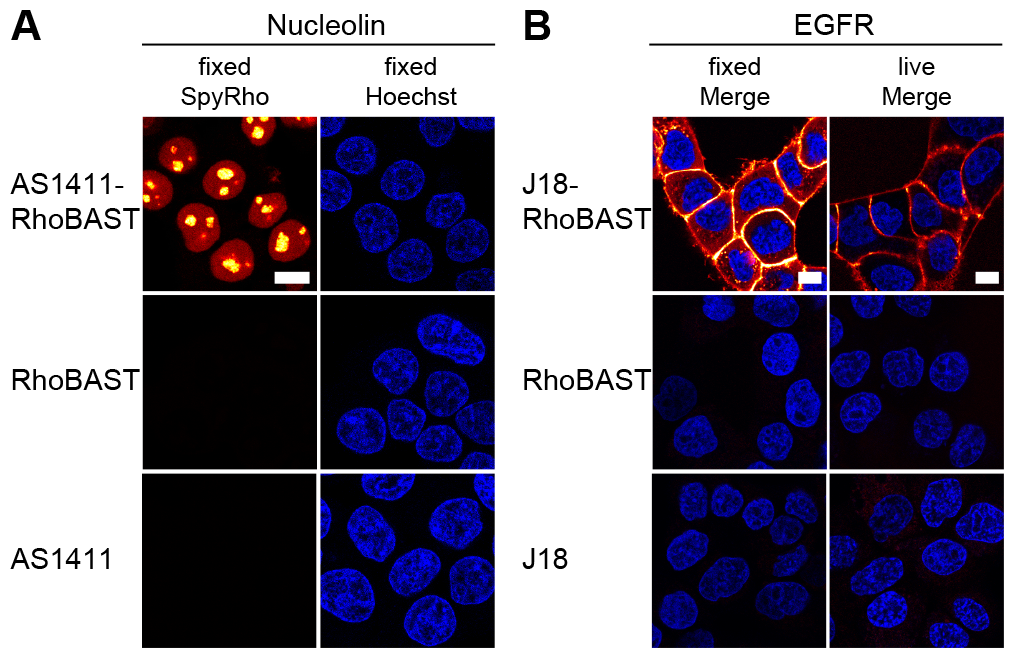
Confocal images of endogenous proteins visualized by bifunctional aptamers in fixed and on live mammalian cells. **A)** Nucleolin was visualized in fixed and permeabilized HEK293T cells using AS1411-RhoBAST (500 nM) and SpyRho (100 nM). **B)** EGFR was visualized in fixed and on live A431 cells using J18-RhoBAST (500 nM) and SpyRho (100 nM). As negative controls, protein-binding aptamer (500 nM) or RhoBAST (500 nM) in the presence of SpyRho are shown (100 nM). Nuclei were stained with Hoechst 33342. Scale bars, 10 µm.

We also targeted the epidermal growth factor receptor (EGFR) using the EGFR-binding RNA aptamer J18.^[28, 29]^ A simple linear fusion of J18 to RhoBAST enabled targeting of endogenous EGFR of fixed as well as live A431 cells (Fig. 2B, Fig. S4). A431 cells are well known for their high expression of EGFR, whereas HeLa cells served as negative controls (Fig. S4D). In this case, the *in vitro* synthesized bifunctional aptamer J18-RhoBAST and SpyRho could be added simultaneously without the need for an additional washing step. Incubation of fixed or live A431 cells with J18-RhoBAST:SpyRho permitted imaging after only 10 minutes of incubation (Fig. 2B, Fig. S4D). The specificity of the bifunctional aptamers was further highlighted by transient transfection of A431 cells with a plasmid encoding a GFP-tagged EGFR and visualization either with AP3-RhoBAST or J18-RhoBAST (Fig. S4C). As expected, EGFR was stained in all A431 cells by J18-RhoBAST and only in GFP-positive cells by AP3-RhoBAST (Fig. S4C).

### Genetically encodable bifunctional RhoBAST aptamers for live-cell protein imaging

A significant advantage of unmodified RNA aptamers lies in their genetic encodability. Therefore, they can be expressed in live cells using various RNA expression systems.^[20, 30, 31]^ SpyRho’s cell-permeability and non-toxic nature enhance its applicability to live-cell experiments.^[22]^ Thus, when combining RhoBAST with another genetically encoded RNA aptamer, such as the GFP-binding AP3 aptamer, the bifunctional aptamer can also be expressed in cells and utilized for imaging proteins in live cells. To this end, we employed the Tornado expression system^[20]^ to express circular AP3-RhoBAST in mammalian cells. We transiently co-transfected COS7 with a plasmid encoding for the GFP-tagged POI and a second plasmid encoding AP3-RhoBAST. This approach enabled the visualization of various subcellular structures, including microtubules using GFP-tagged Tau, the endoplasmic reticulum membrane with GFP-tagged Sec61β, and the outer mitochondrial membrane with GFP-tagged TOMM20 in living cells (Fig. 3). The colocalization of GFP and SpyRho fluorescence indicates the successful imaging of the subcellular structures by the AP3-RhoBAST bifunctional aptamer (Fig. 3). In control experiments, we expressed the circular individual aptamers AP3 and RhoBAST. Cells expressing GFP-tagged proteins and circular RhoBAST displayed a homogeneously distributed cytosolic SpyRho signal, which was not colocalized with GFP fluorescence (Fig. S5). On the other hand, cells expressing GFP-tagged proteins and circular AP3 showed minimal SpyRho fluorescence background and no colocalization with GFP fluorescence (Fig. S5).

**Figure 3:**
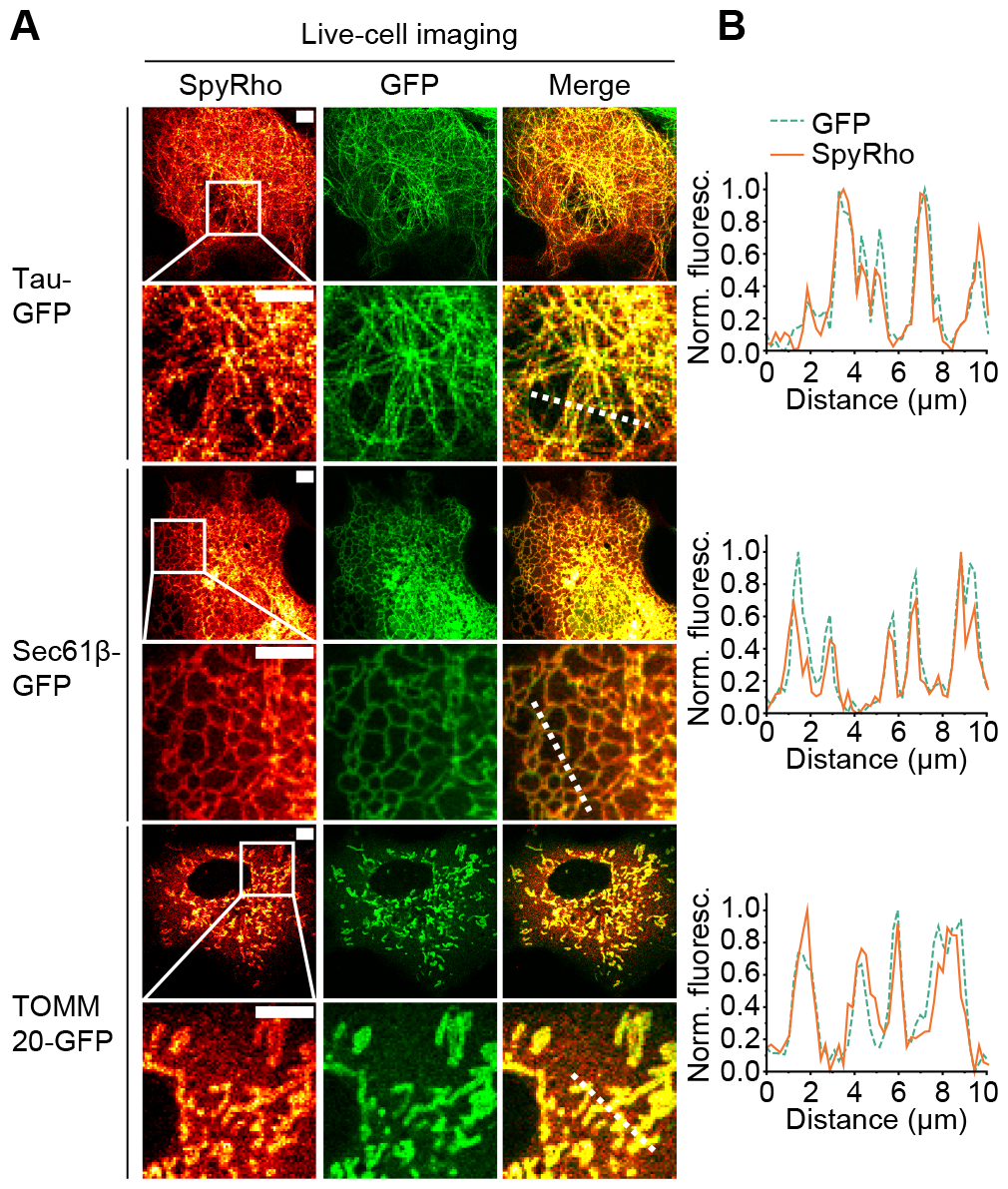
Protein visualization by genetically encoded bifunctional AP3-RhoBAST in live COS7 cells. **A)** Confocal images of live COS7 cells co-expressing Tau-GFP, Sec61β-GFP, or TOMM-20-GFP and circular AP3-RhoBAST. Cells were incubated with SpyRho (100 nM) for 30 minutes before imaging. The white frames indicate the regions of the shown zoom-ins. Scale bars, 5 µm. **B)** Normalized fluorescence profiles of SpyRho (orange, solid line) and GFP (green, dashed line) fluorescence along the dashed lines depicted in A).

### Dual-color protein imaging using bifunctional RhoBAST and SiRA aptamers

Our approach to protein labeling with bifunctional aptamers is modular, i.e., not only the protein-binding aptamer but also the FLAP is exchangeable. To facilitate multiplexing, we introduced silicon rhodamine-binding aptamer (SiRA) as a far-red FLAP with an excellent quantum yield (QY = 0.98) to the bifunctional aptamer system (Fig. 4A).^[32]^ SiRA was connected to AP3 following the same scaffold design and characterization strategy as for AP3-RhoBAST. Consequently, SiRA and AP3 were combined in three constructs using: i) the F30-scaffold, ii) the tetraloop of SiRA, and iii) a 10-nucleotide linker. SiRA has two tetraloops and we examined both positions for the introduction of AP3 (Fig. S6D). Evaluating the fluorescence turn-on and affinity of AP3-SiRA connected via the first loop (UGAA) or via the second loop (UUCG) with the fluorogenic silicon rhodamine (SiR) dye revealed that only fusion into the second loop retained SiRA’s binding and turn-on (Fig. S6B, D, E). Two AP3-SiRA scaffolds exhibited strong binding affinities to SiR with *K*D values of 360 nM (loop) and 417 nM (F30-scaffold), while a larger *K*D of 1047 nM was observed for the scaffold with flexible linker (Fig. 4B, Fig. S6B, C). The highest fluorescence turn-on was measured for AP3-SiRA in the F30-scaffold (7-fold, Table S1). For the *in cellulo* investigations of AP3-SiRA, we chose the same procedure for H2B-GFP visualization in fixed and permeabilized HEK293T cells, as described above. Staining with each AP3-SiRA construct consistently resulted in colocalization of the GFP and SiR fluorescence signals for all fusion designs, whereas minimal fluorescence was observed for the negative control samples SiRA and AP3 (Fig. 4, Fig. S6D). It is important to note that the fluorescence turn-on of AP3-SiRA varied significantly among the three fusion designs (Figure 4C, D, Table S1). We observed 7-, 4-, and 2-fold turn-on values for F30-, loop- and linker-scaffolds, respectively. Since AP3-SiRA in the F30-scaffold displayed the most favorable *in cellulo* performance, it was selected for all further experiments.

**Figure 4:**
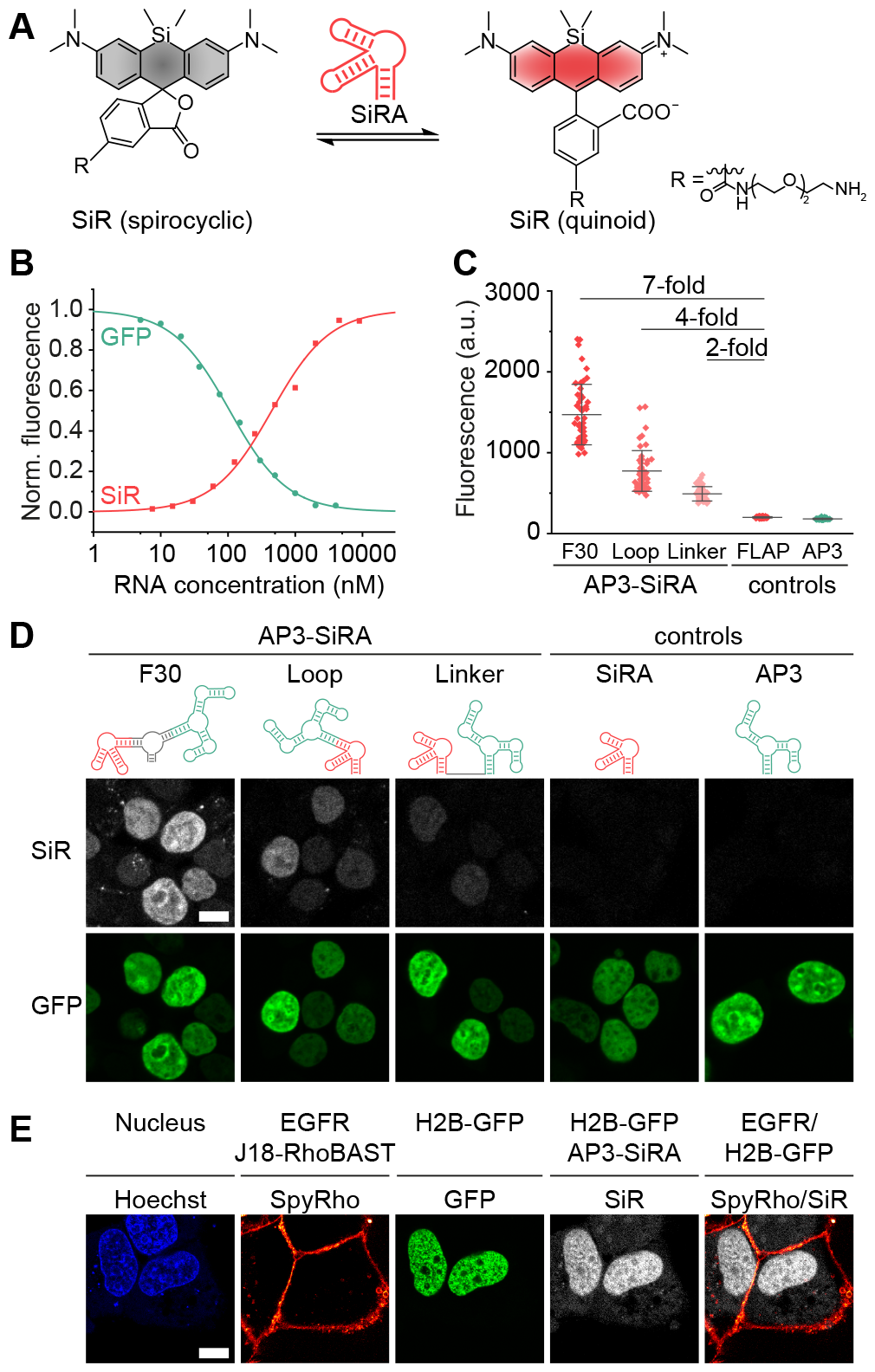
Characterization of AP3-SiRA and its application in dual-color protein imaging. **A)** Equilibrium of silicon rhodamine (SiR) between the non-fluorescent spirocyclic form and the fluorescent quinoid form. SiR lights up upon binding to the aptamer, SiRA. **B)** Binding isotherms of AP3-SiRA (F30-scaffold) with SiR (10 nM, red line) and GFP (5 nM, green line) giving dissociation constants of *K*_D_ = 417 ± 29 nM and *K*_D_ = 105 ± 18 nM, respectively. Minimal and maximal fluorescence intensities were normalized to 0 and 1, respectively. *K*_D_-values are given as mean ± s.d., *N* = 3 independent experiments. **C)** Quantification of SiR fluorescence intensities (mean ± s.d., *N* = 50 cells) of HEK293T cells expressing H2B-GFP from confocal images as shown in D). **D)** Confocal images of fixed and permeabilized HEK293T cells expressing H2B-GFP. Cells were incubated with 1000 nM aptamer, washed and stained with 500 nM SiR. Scale bar, 10 µm. **E)** Confocal images of dual-color protein labeling. EGFR was visualized with J18-RhoBAST (500 nM) and SpyRho (100 nM)) in live A431 cells expressing H2B-GFP. Then, cells were fixed, permeabilized, and H2B-GFP was visualized using AP3-SiRA (1000 nM) and SiR (500 nM). Scale bar, 10 µm.

Utilizing the bifunctional aptamers J18-RhoBAST and AP3-SiRA, we successfully visualized EGFR and H2B-GFP within the same A431 cells. For this purpose, A431 cells were transiently transfected to express H2B-GFP. Since EGFR can be visualized in live cells, we adopted a sequential procedure for this experiment. First, the transfected live A431 cells were stained with J18-RhoBAST:SypRho to visualize EGFR. Subsequently, the cells were washed, fixed, and permeabilized. After washing, the cells were incubated with AP3-SiRA and stained with SiR to achieve the desired visualization of histone H2B (Fig. 2 D). All aptamers used in this experiment exhibited high specificity towards their respective targets as confirmed through multiple negative control experiments, including all aptamers individually (Fig. S7).

### Super-resolved protein imaging using RhoBAST-PAINT

RhoBAST:SpyRho has already proven to be an excellent tool for SMLM-based super-resolved RNA imaging, operating on the same principles as DNA-PAINT.^[22]^ Here, we advance its capabilities to visualize proteins with super-resolution using bifunctional aptamers, and name this technology RhoBAST-PAINT. To validate the feasibility of this approach, we first determined the kinetic rate coefficients of the bifunctional AP3-RhoBAST and SpyRho using stopped-flow kinetic measurements. Titrating SpyRho with different AP3-RhoBAST concentrations and measuring the fluorescence intensity increase over time revealed an association rate coefficient (*k*a) of 3.8 × 10^7^ M^-1^s^-1^ and a dissociation rate coefficient (*k*d) of 1.2 s^-1^ (Fig. S8, Table S). These values were comparable to those of the RhoBAST:SpyRho complex (*k*a = 2.1 × 10^7^ M^-1^s^-1^ and *k*d = 1.8 s^-1^),^[22]^ further supporting the applicability of bifunctional aptamers in RhoBAST-PAINT. Next, to test the performance of bifunctional aptamers in super-resolved protein visualization, the GFP-tagged histone H2B was imaged with AP3-RhoBAST and SpyRho in fixed HEK293T cells. To minimize multiple simultaneous blinking events in the same area, we used a low concentration of SpyRho (1 nM). Reconstruction of 40,000 frames resulted in a super-resolved image with a localization precision of 27 nm for H2B-GFP and a resolution of 69 nm obtained by decorrelation analysis (Fig. 5A–C).^[33]^ Subsequently, we imaged the endogenous EGFR in fixed A431 cells using J18-RhoBAST and SpyRho with the RhoBAST-PAINT approach achieving a comparable localization precision and resolution (26 nm, 67 nm, respectively. Fig. 5D–F).

**Figure 5:**
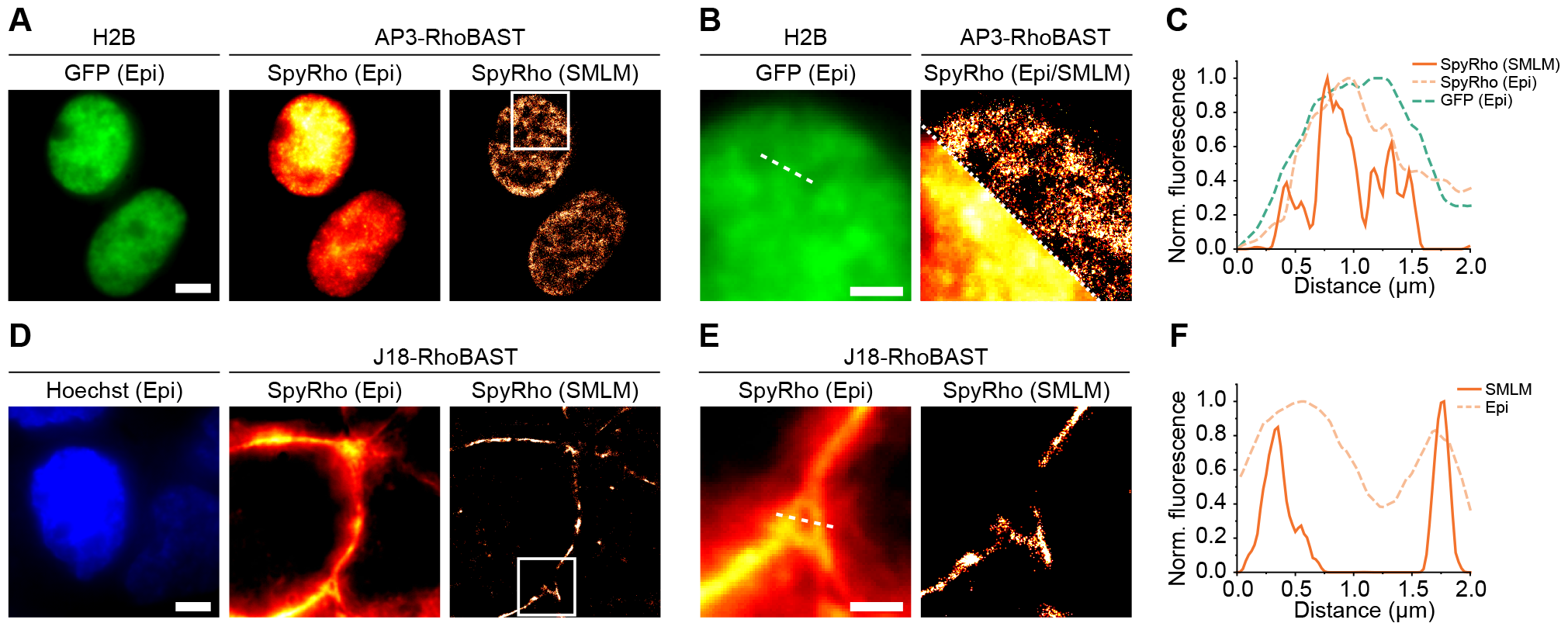
Application of bifunctional aptamers in RhoBAST-PAINT. **A)** Fixed and permeabilized HEK293T cells expressing H2B-GFP were stained with AP3-RhoBAST (500 nM) and SpyRho (1 nM). SMLM images were reconstructed from 40,000 frames with an exposure time of 30 ms. Scale bar, 5 µm. **B)** Zoom-in of the region indicated by the white frame in A). Scale bar, 2 µm. **C)** Normalized fluorescence intensity profiles of epifluorescence (dashed lines) and super-resolved (solid line) images along the dashed line shown in B). **D)** Visualization of EGFR in fixed A431 cells stained with J18-RhoBAST (500 nM) and SpyRho (1 nM). SMLM images were reconstructed from 3000 frames. Scale bar, 5 µm. **E)** Zoom-in of the region indicated by the white frame in D). Scale bar, 2 µm. F) Normalized fluorescence intensity profiles of epifluorescence (dashed lines) and super-resolved (solid line) images along the dashed line shown in E).

### Comparative analysis of bifunctional RhoBAST aptamers and antibodies in protein imaging via CLSM and SMLM

Focusing on the preparative handling and the performance in both confocal and single-molecule localization microscopy, we compared the bifunctional aptamer to the commonly used immunofluorescence antibody approach. Therefore, we visualized Synaptophysin-GFP, EGFR, and nucleolin using either bifunctional RhoBAST aptamer or a combination of primary and secondary antibodies. For a fair comparison, we used a secondary antibody conjugated with the tetramethylrhodamine fluorophore, which is structurally similar to SpyRho and can be excited with the same laser wavelength. For visualization with bifunctional aptamers, the above-described protocols were applied, while for antibody staining, we adhered to the manufacturers’ protocols. Traditional immunostaining involves a lengthy protocol, including BSA blocking, multiple washes, and incubation with primary and secondary antibodies. Our approach, utilizing *in vitro* synthesized bifunctional aptamers, simplifies the process significantly, requiring only one wash, or in case of EGFR visualization no washes at all. Synaptophysin-GFP in fixed neurons was visualized with AP3-RhoBAST and SpyRho, or with anti-synaptophysin primary antibody and secondary rhodamine-labeled antibody, resulting in qualitatively comparable structures and colocalization with GFP fluorescence (Fig. 6A, Fig. S9). For EGFR targeting, similar localizations were observed using either J18-RhoBAST:SpyRho or anti-EGFR antibody and secondary rhodamine-labeled antibody in fixed A431 cells (Fig. 6A). Lastly, in targeting nucleolin in fixed HEK293T cells, we used AS1411-RhoBAST and SpyRho or anti-nucleolin antibody with a secondary rhodamine-labeled antibody (Fig. 6A). The aptamer approach exhibited superior bleaching resistance: fluorescence intensity decreased by only 13% in multiple frames (# frames = 23), compared to 39% with antibodies. This enhanced performance is attributed to fast dye exchange and increased photostability of SpyRho upon binding to RhoBAST.

**Figure 6:**
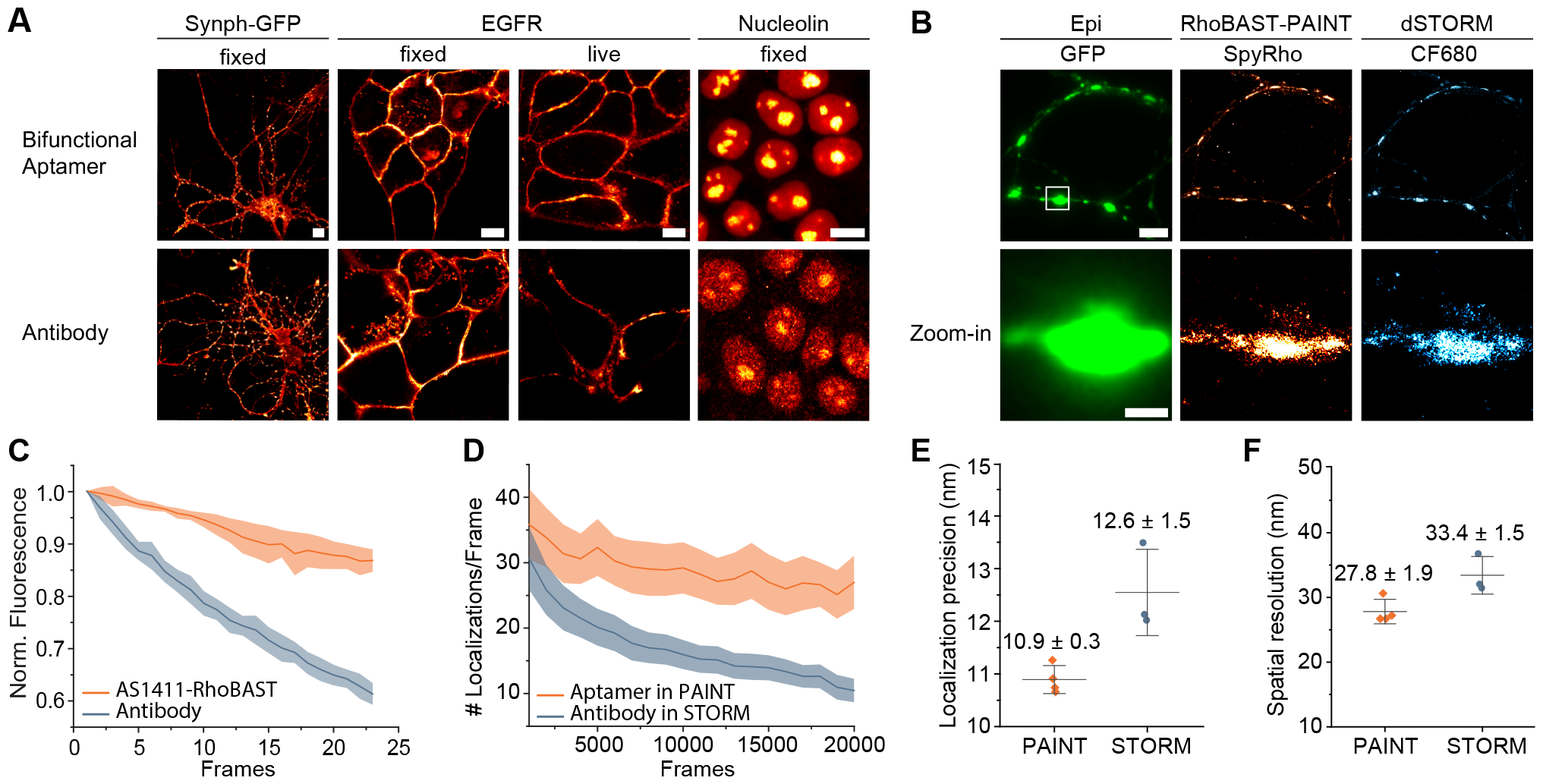
Comparison of bifunctional aptamers to antibodies in CLSM and SMLM. **A)** Confocal images comparing staining with protein-binding-aptamer-RhoBAST and SpyRho to immunostaining with primary and rhodamine-labeled secondary antibodies. From left to right panel: Synaptophysin-GFP (Synph-GFP) in fixed and permeabilized hippocampal cells was visualized either with AP3-RhoBAST:SpyRho or anti-synaptophysin primary and rhodamine-labeled secondary antibodies. EGFR in fixed or live A431 cells was visualized either with J18-RhoBAST:SpyRho or anti-EGFR primary and rhodamine-labeled secondary antibodies. Nucleolin in fixed and permeabilized HEK293T was visualized either with AS1411-RhoBAST:SpyRho or anti-nucleolin primary and rhodamine-labeled secondary antibodies. Scale bars, 10 µm. **B)** Comparison of RhoBAST-PAINT and *d*STORM in hippocampal cells expressing synaptophysin-GFP. Cells were fixed, permeabilized and incubated with AP3-RhoBAST (500 nM), washed and imaged using SpyRho (1 nM). Thereafter, the sample was washed and immunostained using anti-synaptophysin primary antibody and CF680-labeled secondary antibody. Reconstruction of 20,000 frames each yielded the super-resolved RhoBAST-PAINT and *d*STORM image, respectively. Scale bars, 5 µm for upper panels and 1 µm for zoom-in. **C)** Normalized fluorescence decrease of aptamer or antibody staining of nucleolin in fixed and permeabilized HEK293T cells shown in A (fourth panel) over 23 frames confocal imaging using a laser power of 440 µW (mean ± s.d., *N =* 25 nucleolin regions of interests). **D)** Decrease of the number of localizations per frame over time (30 ms per frame) of the experiment shown in B). **E)** Nearest neighbor analysis of *d*STORM or RhoBAST-PAINT images, as shown in B), gave the localization precision. Data represent mean ± s.d. (*N* = 3 for *d*STORM, *N* = 4 for RhoBAST-PAINT). **F)** Spatial resolution of the SMLM images achieved by *d*STORM or RhoBAST-PAINT, as shown in B), were determined by decorrelation analysis. Data represent mean ± s.d. (*N* = 3 for *d*STORM, *N* = 4 for RhoBAST-PAINT).

*d*STORM is a widely used method for super-resolved imaging of proteins, in particular for immunofluorescence applications.^[1, 34, 35]^ Therefore, we next compared RhoBAST-PAINT and *d*STORM within the same sample, using the previously described maS^3^TORM setup.^[34]^ In *d*STORM, photoswitchable fluorophores capable of transitioning between a fluorescent and non-fluorescent state are used to generate the required blinking. Rat hippocampal cells expressing GFP-tagged synaptophysin were fixed and permeabilized. To ensure minimal interference between aptamer- and antibody-based methods, we utilized GFP-binding AP3-RhoBAST, while an anti-synaptophysin antibody directly targeted synaptophysin. Furthermore, fluorophores with different excitation and emission wavelengths were chosen, namely SpyRho and the established *d*STORM dye CF680.^[34, 36]^ First, super-resolved images of synaptophysin-GFP were obtained by using AP3-RhoBAST with SpyRho (Fig. 6B, Supplementary Movie). This experiment yielded a localization precision of 10.9 nm according to nearest neighbor analysis^[37]^ and a resolution of 27.8 nm, as quantified by decorrelation analysis (Fig. 6E, F).^[33]^ After washing the cells, *d*STORM images were acquired using an anti-synaptophysin primary antibody and CF680-labeled secondary antibody (Fig. 6B, Supplementary Movie). Analysis of *d*STORM data resulted in a slightly lower resolution of 33.4 nm with a localization precision of 12.6 nm (Fig. 6E, F). The comparable resolutions of *d*STORM and RhoBAST-PAINT proved the applicability of bifunctional aptamers in super-resolved protein imaging. Moreover, the quantitative analysis of the number of identified localizations over time revealed substantial decrease of 66% in *d*STORM, compared to only 25% decrease in RhoBAST-PAINT (Fig. 6D).

## Conclusion

In summary, we have introduced bifunctional aptamers for protein visualization, leveraging the exceptional photophysical properties of fluorescent light-up aptamers and the specificity of protein-binding aptamers. This approach offers versatility and adaptability on both sides, the FLAP and the protein-binding aptamer. FLAPs are available over a broad spectral range as well as with varying photophysical properties, and several thousand protein-binding aptamers are known.^[38, 39]^ The demonstrated combination of RNA/DNA aptamer fusions further extends the palette of possible target proteins. Notably, RNA aptamers are easily synthesized *in vitro* and cost-effective in comparison to antibodies, with the added advantage of genetic encodability, enabling their application in living cells. While transient transfection may introduce challenges due to varying expression levels of proteins and bifunctional aptamers, AP3-RhoBAST has demonstrated utility. It is essential to recognize that, unlike some protein-binding aptamers,^[12]^ AP3-RhoBAST’s interaction with GFP minimizes changes or inhibition of the target protein’s function, since the aptamer binds to the GFP tag rather than directly to the target protein itself. Based on bifunctional aptamers, we developed RhoBAST-PAINT, successfully utilizing the RhoBAST:SpyRho system for super-resolution imaging of proteins due to excellent photophysical properties and fast association and dissociation kinetics. Here, proteins were imaged on the nanometer scale yielding 28–69 nm resolution depending on the imaging depth and the sample. The bifunctional aptamers are approximately 7-fold smaller in weight and size than conjugates of primary and secondary antibodies, so that the expected linkage error to the target is smaller.^[1]^ The smaller tag size may also provide better access to challenging epitopes and improved tissue penetration. Furthermore, in the RhoBAST:SpyRho system, each RhoBAST binds a single fluorogen, providing single-molecule blinking in PAINT, whereas an antibody is usually conjugated to several fluorophore molecules with a heterogeneous distribution. RhoBAST’s non-covalent interaction with SpyRho allows for exchange of photobleached fluorophores, enhancing localization precision and resolution. The concept of bifunctional aptamers for protein visualization holds promise for future applications in advanced fluorescence microscopy and single-molecule localization-based super-resolution imaging.

## Supporting information

Supplementary Movie_RhoBAST-PAINT_(left)_dSTORM_(right)

## Author contributions

F.G., B.B., A.J., G.U.N., T.K., M.S. designed the study. F.G. and N.v.d.B performed all *in vitro* and *in cellulo* experiments and confocal microscopy imaging. F.G. cloned plasmids. F.G. and M.S.V. performed and analyzed SMLM experiments in mammalian cells. F.G. and M.K. performed and analyzed SMLM and *d*STORM experiments in hippocampal cells. F.G. wrote the manuscript and all authors participated in revising and editing.

## Acknowledgments

The authors are grateful to D. Englert providing plasmids coding for GFP-tagged proteins and to K. Nienhaus supporting SMLM imaging. We acknowledge the Nikon Imaging Center, Heidelberg giving access to their microscopy facility. We thank M. Mayer and L. Rohland for assistance with stopped-flow experiments. We thank M. Schmitt and F. Gleiche for excellent technical assistance in hippocampal cell culture.

