## Supporting Information for "Super-resolved protein imaging using bifunctional light-up aptamers"

[a]Institute of Pharmacy and Molecular Biotechnology (IPMB), Heidelberg University, Heidelberg, Germany. [b]Department of Nuclear Medicine, Heidelberg University Hospital, Heidelberg, Germany. [c]Department of Functional Neuroanatomy, Institute for Anatomy and Cell Biology, Heidelberg University, Heidelberg, Germany. [d]Institute of Applied Physics (APH), Karlsruhe Institute of Technology (KIT), Karlsruhe, Germany. [e]Department of Chemical Biology, Max Planck Institute for Medical Research, Heidelberg, Germany. [f]Institute of Nanotechnology (INT), Karlsruhe Institute of Technology (KIT), Eggenstein-Leopoldshafen, Germany. [g]Institute of Biological and Chemical Systems (IBCS), Karlsruhe Institute of Technology (KIT), Eggenstein-Leopoldshafen, Germany. [h]Department of Physics, University of Illinois at Urbana−Champaign, Urbana, IL, USA.

***** To whom correspondence should be addressed.

### Experimental methods

#### General

Reagents were purchased from Thermo Fisher Scientific or Sigma-Aldrich and used without further purification. DNA oligonucleotides were purchased from Integrated DNA Technologies. All experiments were carried out using deionized water purified using a Milli-Q Synthesis A10 Water Purification system or EQ 7000 system (Merck). Absorbance measurements were conducted using a Cary50 UV/Vis spectrometer (Varian). Fluorescence measurements were recorded on a FP-8500 fluorospectrometer (JASCO) at 25°C. Stopped-flow experiments were carried out using a SX-18M stopped-flow spectrometer (Applied Photophysics). Fluorophores (SiR,[1] SpyRho,[2] GFP (addgene plasmid #29663)) were synthesized lab stocks. SpyRho was also purchased from Spirochrome. Prediction of RNA folding were made using mfold[3] and visualized by VARNA software.[4]

#### *In vitro* synthesis of RNA aptamers

*In vitro* transcription was conducted using T7 RNA polymerase (lab stock), 1 µM dsDNA template, 2 mM NTP, 10 mM DTT in transcription buffer (40 mM Tris-HCl (pH 8.1), 1 mM Spermidine, 22 mM $MgCl_2$, 0.01% Triton X-100, 5% DMSO, 10 mM DTT) over 4 h at 37°C. Thereafter, the reaction was incubated with DNaseI (10 U/µL) for 30 min at 37°C. The RNA was purified by denaturing polyacrylamide gel electrophoresis (10%). The desired band was visualized by UV-shadowing, excised, crushed and eluted in 0.3 M NaOAc buffer (pH 5.5). The product was obtained by isopropanol precipitation.

#### *In vitro* synthesis of RNA-DNA conjugated aptamers

The *in vitro* synthesized RNA (1 µM) was ligated to the single-stranded DNA starting with adenosine monophosphate (rA, 10 µM) using T4 RNA Ligase I (0.5 U/µL) and ATP (1 mM) in T4 RNA Ligase Reaction Buffer (New England Biolabs) supplemented with PEG8000 (20% w/v, Jena Bioscience). The product was purified by denaturing polyacrylamide gel electrophoresis (10%) and isopropanol precipitation as stated above.

#### *In vitro* characterization of aptamer:dye complexes

For all *in vitro* measurements and in fixed cell experiments, aptamers were treated as follows to ensure proper folding: RNA or RNA-DNA conjugates were dissolved in water, incubated for 2 min at 75°C and cooled to 25°C at a rate of 5°C/min. Then the desired buffer concentrate (6x) was added and incubated for 10 min. If not stated otherwise, experiments were conducted in aptamer selection buffer, ASB (20 mM HEPES pH 7.4, 1 mM $MgCl_2$, 125 mM KCl) or ASBT (ASB supplemented with 0.05% (v/v) Tween20). For fluorescence measurements, dyes and aptamer:dye complexes were excited at the maximum excitation wavelength and intensities were detected at the maximum emission wavelength. Excitation and emission spectra of fluorophores were recorded in the presence and absence of aptamer. Minimal and maximal fluorescence intensities were normalized to 0 and 1, respectively.

Dissociation constants ($K_D$) were obtained through measuring the fluorescence change of dyes upon binding to the aptamer. A fixed fluorophore concentration (10 nM) was titrated with different aptamer concentrations in ASBT. The measured intensities were fitted as previously published.[5] Minimal and maximal fluorescence intensities were normalized to 0 and 1, respectively.

The fluorescence turn-on values represent the ratio of fluorescence intensities with and without aptamers present. The turn-on values of SiRA within bifunctional aptamer constructs were determined using SiR (5 nM) and an excess of aptamer (5 µM). The turn-on values of RhoBAST within bifunctional aptamer constructs were determined using SpyRho (1 µM) and an excess of aptamer (5 µM). The fluorescence turn-off value represents the ratio of fluorescence intensities in the absence and in the presence of aptamer. The turn-off values of AP3 within bifunctional aptamer constructs were measured using GFP (10 nM) and an excess of aptamer (5 µM).

Kinetic rate coefficients of AP3-RhoBAST:SpyRho were obtained using stopped-flow kinetic measurements as described previously. [2, 5] Briefly, SpyRho (5 nM) was titrated with different concentrations of AP3-RhoBAST in ASBT at 25°C and the fluorescence increase over time was measured. The observed kinetic rate coefficients $k_{obs}$ were obtained using equation (1) with $F(t)$ as time-dependent fluorescence, $F_0$ as initial fluorescence:

$$F(t) = (F_0 - F_\infty)e^{-k_{obs}t} + F_\infty \quad (1)$$

By linear fitting of the observed kinetic rates versus aptamer concentration, the association rate coefficient $k_a$ and the dissociation rate coefficient $k_d$ were determined using equation (2):

$$k_{obs} = k_a[\mathrm{RNA}] + k_d \quad (2)$$

At each concentration, the fluorescence increase over time was measured in five technical replicates with the stopped-flow spectrometer. Three independent measurements were performed to determine the kinetic rate coefficients as mean ± s.d.

**DNA cloning and plasmids**

Sequences can be found in Table S4 and S5. Double stranded DNA inserts of AP3, RhoBAST, and AP3-RhoBAST (connected via loop) with NotI and SacII restriction sites were obtained by PCR. The inserts were double digested using NotI and SacII according to the manufacturer´s protocol and cloned into the NotI and SacII double digested pAV-U6+27-Tornado-Broccoli (Addgene, plasmid #124360)[6] to yield pAV-U6+27-Tornado-AP3, pAV-U6+27-Tornado-RhoBAST, pAV-U6+27-Tornado-AP3-RhoBAST. Furthermore the following plasmids were used: pCMV-EGFR-GFP (Addgene, plasmid #32751),[7] pRK5-EGFP-Tau (Addgene, plasmid #46904),[8] pcDNA5-TOMM20-GFP,[2] pcDNA5-H2B-GFP,[2] pcDNA5-Sec61β-GFP,[2] pAM-mGFP (membrane-bound GFP),[9] pAM-Synaptophysin-EGFP.[10]

**Cell culture and chemical fixation**

**Secondary cell culture**. HEK293T, COS7, and A431 cells were cultivated at 5% $CO_2$ and 37°C in DMEM (Dulbecco´s Modified Eagle Medium containing high D-glucose, 25 mM HEPES, L-glutamine, without phenol red) supplemented with fetal bovine serum (FBS, 10% (v/v)), 100 U/mL penicillin and 100 µg/mL streptomycin. Cells were cultured in T25-flasks and split every 2–3 days. For imaging experiments, cells were seeded in µ-slides (glass bottom) from Ibidi with 8 or 18 wells. If HEK293T cells were seeded, slides were coated with poly-D-lysine. For washing the cells, DPBS (Dulbecco´s Phosphate Buffered Saline (modified, without calcium and magnesium chloride)) was used. Cells were fixed by incubating with 4% PFA for 10 min. Then cells were incubated with GlyPBS (DPBS supplemented with 1 mM $MgCl_2$ and 100 mM glycine) for 5 min and washed twice with MgPBS (DPBS supplemented with 1 mM $MgCl_2$). If needed, cells were permeabilized using 0.1% Triton X-100 in MgPBS for 15 min and afterwards washed three times with MgPBS. Live cell experiments were carried out in warm L15 medium (Leibovitz´s medium).

**Primary cell culture.** Primary hippocampal cell cultures were established from E19 Sprague- Dawley rats (Charles River Laboratories). Pregnant rat dams were anesthetized, and embryos were euthanized in compliance with the German animal welfare guidelines (protocol T-33/21 approved by the Regierungspraesidium Karlsruhe). Hippocampi were dissected from brains of E19 embryos in ice-cold DPBS under a dissecting microscope and then collected in cold HBSS (Hank's balanced salt solution, Invitrogen). Subsequently, hippocampi were incubated in 0.25% trypsin in HBSS for 20 min at 37°C. Enzymatic activity of trypsin was stopped by addition of warm DMEM (high glucose, L-glutamine, sodium pyruvate, phenol red) supplemented by 10% FBS, 100 U/mL penicillin/streptomycin and 2 mM L-glutamine to the cell suspension at a 2:1 ratio. After five minutes, the supernatant was removed and cells were washed with DMEM. Cells were mechanically dissociated and subsequently filtered through a cell strainer with a mesh width of 100 µm (Greiner). Hippocampal cells were collected by centrifugation at 500 g for 5 min and resuspended in DMEM supplemented with 10% FBS, 100 U/mL penicillin/streptomycin, and 2 mM L-glutamine. Hippocampal cells were seeded on poly-L-lysine-coated glass-bottom dishes (µ-slides glass bottom, Ibidi or P35G-0-14-C, MatTek). Hippocampal cultures were stored in an incubator maintaining 37°C and 5% $CO_2$. Medium was exchanged by neurobasal medium (21103-049, Invitrogen) with 1x B27 (Gibco), 2 mM L-glutamine, and 100 U/mL penicillin/streptomycin after 5 hours. After 14 days in the incubator, hippocampal cultures were transduced using adeno-associated virus (AAV) with a pAM vector plasmid carrying a synaptophysin-EGFP fusion construct or membrane-bound GFP (mGFP) with a MARCKS protein-derived membrane tag containing palmitoylation sites. Five days after transduction, hippocampal neurons were rinsed twice with cytoskeletal buffer (160 mM 1,4-piperazinediethanesulfonic acid pH 6.8, 10 mM EGTA, 4 mM $MgCl_2$) at 37°C and chemically fixed using 4% PFA in cytoskeletal buffer supplemented by 4% sucrose at 37°C for 10 min. If needed, hippocampal neurons were permeabilized using 0.1% Triton X-100 in PBS for 10 min and blocked in 5% fetal calf serum (FCS) with 100 mM glycine for 15 min.

**Confocal microscopy**

Confocal laser scanning microscopy was performed at a point scanning Nikon A1R confocal microscope based on a fully inverted Nikon Ti2 equipped with a hybrid scanner (galvo/resonant), four-channel detection and a live-cell chamber for controlled temperature and humidity. A Nikon N Apo 60× NA 1.4 λs OI (WD 0.14 mm, FOV 0.21 × 0.21 mm) or a Nikon Plan Fluor 40× NA 1.3 λs OI (WD 0.2 mm, FOV 0.32 × 0.32 mm) were used as objectives. Hoechst 33342 (100 µg/mL) was used as nuclear stain. Hoechst was excited at 405 nm and the emission was detected via a 450 ± 25 nm filter set. GFP was excited at 488 nm and the emission was detected via a 525 ± 25 nm filter set. SpyRho/TMR was excited at 561 nm and the emission was detected via a 595 ± 50 nm filter set. SiR was excited at 640 nm and the emission was detected via a 700 ± 37.5 nm filter set. Images were recorded using the Nikon Elements software and processed with Fiji.[11] For quantification, cell and nuclear segmentation was performed using cellpose.[12]

**Fixed cell confocal microscopy.** Prior to imaging, cells were washed with ASB, incubated for 30 min with *in vitro* synthesized, freshly folded aptamer and washed once with ASB. Then dye was added, and cells were imaged after 15 min of incubation at 25°C. For visualizing EGFR, aptamer and dye were premixed and added without further washing steps. Primary and secondary antibodies (Table S6) were used according to the supplier’s protocols.

**Live-cell confocal microscopy.** Seeded HEK293T, A431, or COS7 live cells were transiently transfected using FuGENE HD (Promega) 48 h prior to imaging. For co-transfection, plasmids coding for GFP-tagged proteins and pAV-U6+27-Tornado-aptamer were mixed in a 1:10 ratio. Dual-color imaging was conducted sequentially. First, EGFR in living A431 cells expressing H2B-GFP was imaged using J18-RhoBAST (500 nM) and SpyRho (100 nM), then the cells were fixed and permeabilized. Secondly, H2B-GFP was visualized by incubating the cells with AP3-SiRA (loop, 1000 nM), washing them once, and adding SiR (500 nM).

**Single-molecule localization microscopy (SMLM)**

**RhoBAST-PAINT in HEK293T and A431 cells**. HEK293T cells expressing H2B-GFP were fixed, permeabilized, and incubated with bifunctional aptamer (500 nM) for 30 min. Then, cells were washed once with ASB, incubated with SpyRho (1 nM) prior to imaging. A431 cells were fixed and incubated with J18-RhoBAST:SpyRho (500 nM, 1 nM) for 30 min prior to imaging. SMLM was conducted at a homebuilt widefield microscope based on an Axio Observer Z1 frame (Zeiss) with a quad-band dichroic mirror (z 405/473/561/640, AHF) and an Ixon Ultra X-7759 EMCCD camera (Andor) with a pixel size of 109 × 109 $nm^2$. Lasers with a wavelength of 561 nm (Gem 561, Laser Quantum) and 473 nm (Gem 473, Laser Quantum) were applied using dichroic mirrors and an acousto-optical tunable filter (AOTFnC-400.650, A-A Opto-Electronic). To expand the laser beam, two achromatic lenses (focal lengths 10 mm, 100 mm, Thorlabs) were used. GFP was excited with 473 nm (laser power 200 µW) and SpyRho was excited with 561 nm (laser power 20 mW). Acquisitions were made approximately 2 µm above the glass coverslip. H2B-GFP images were reconstructed from 40,000 frames with an exposure time of 30 ms. For EGFR visualization, 3000 frames with an exposure time of 100 ms were collected. For epifluorescence images, usually 300 frames were averaged using Fiji software.[11] SMLM analysis was performed as described previously.[5] In brief, SMLM images were reconstructed by using custom-written a-livePALM software, which is based on Gaussian fitting with the maximum likelihood estimation algorithm to determine positions of fluorescence emitters and their uncertainties.[13] Stage drift was corrected via cross-correlation analysis.[14] Spatial resolution was determined through decorrelation analysis.[15]

**RhoBAST-PAINT and immunostaining-based *d*STORM in hippocampal neurons.** Super-resolved imaging in hippocampal neurons was carried out at the maS$^3$TORM setup developed by Klevanski *et al.*[16] To compare *d*STORM and RhoBAST-PAINT performance within the same sample, hippocampal cells transiently expressing Synaptophysin-EGFP were first subjected to aptamer staining. Here, cells were incubated with AP3-RhoBAST (500 nM) for 30 min, washed with ASB, and stained with SpyRho (1 nM). For subsequent image alignment, fluorescent TetraSpeck microspheres (T7279, Invitrogen) were applied to cell samples for 10 min followed by two brief washing steps. RhoBAST-PAINT imaging of multiple regions of interest was conducted using the 561 nm laser at a power of ~0.12 kW/cm² for 20,000 frames with an exposure time of 30 ms. Subsequently, the sample was washed and subjected to immunohistochemistry for the *d*STORM procedure. To that end, neurons were incubated with the primary antibody against synaptophysin (Table S6) in 0.5% FCS, washed three times with PBS for a total of 15 min, and incubated with the secondary antibody conjugated to CF680 fluorophore at 25°C (Table S6). After three washing steps, *d*STORM imaging buffer (100 mM β-mercaptoethylamine (MEA) in PBS, 15 mM KOH, pH 8) was applied to the sample. *d*STORM acquisition of the regions previously imaged in the RhoBAST-PAINT mode was conducted using the 660 nm laser at ~2.6 kW/cm² for 20,000 frames with an exposure time of 30 ms. The focal plane was maintained within 500 nm of the glass surface. Images were reconstructed using the rapidSTORM[17] algorithm with local relative thresholding. Individual localizations were fitted followed by post-processing using an in-house written software for drift correction, linking of blinking events that remained in the 'on' state in consecutive frames, and image rendering.[16] Localization precision was calculated by nearest neighbor analysis using LAMA-software.[18] This analysis was conducted on data from a fiducial-free area of approximately $19^2$ µm² and 5000 frames, where localizations were not connected across frames. Spatial resolution was assessed through decorrelation analysis[15] on reconstructed fiducial-free SMLM image areas that were rendered based on intensity counts and by applying σ obtained by nearest neighbor analysis.

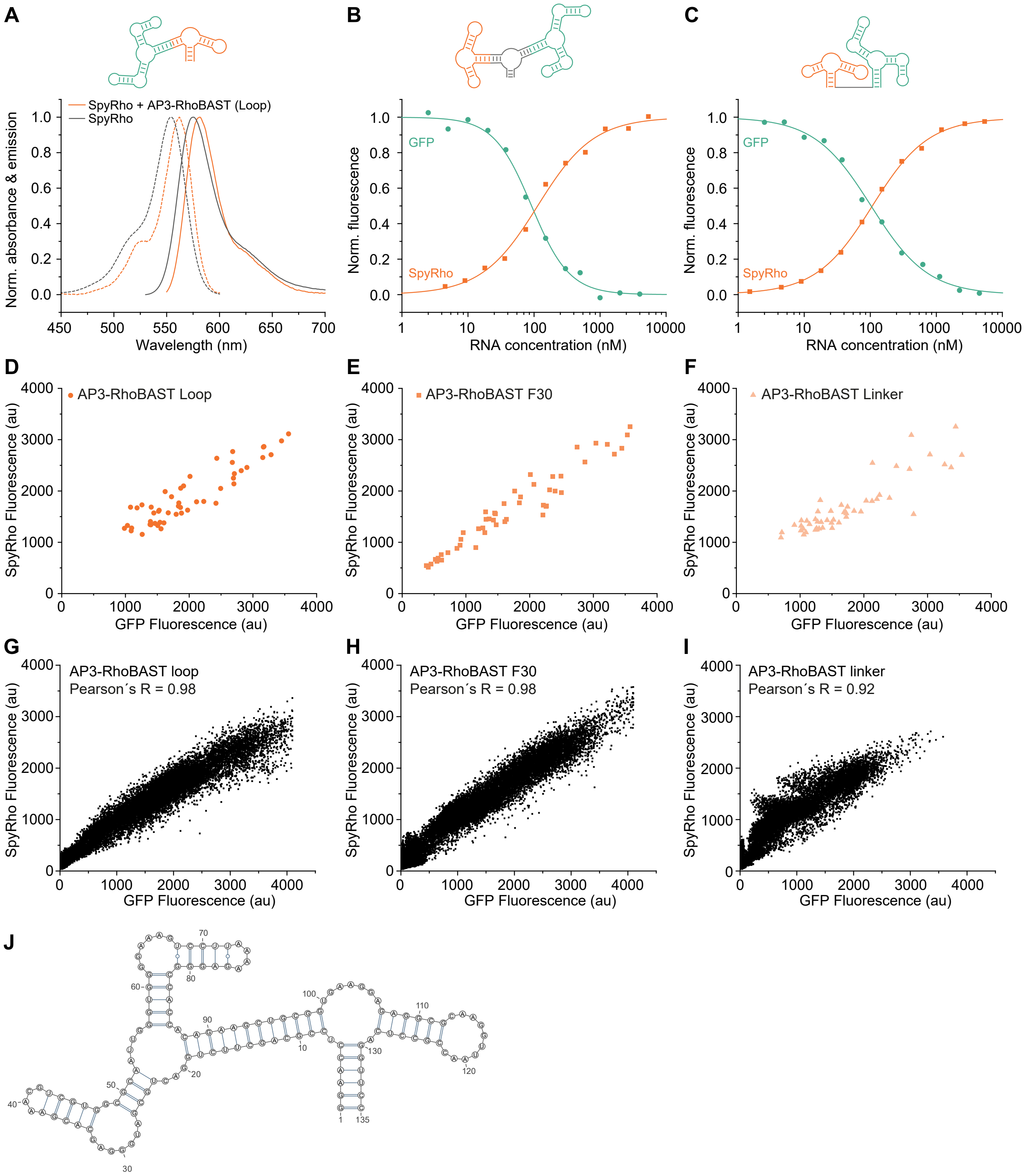


**Supplementary Figure S1: Comparison of different fusion constructs of bifunctional aptamer AP3-RhoBAST. A)** Normalized absorbance (dotted lines) and emission (solid lines) spectra of SpyRho in the presence and absence of AP3-RhoBAST (5400 nM, loop). **B)** Binding isotherms of AP3-RhoBAST (F30) with SpyRho (10 nM, orange line) and GFP (5 nM, green line) giving dissociation constants of $K_D$ = 122 ± 10 nM and $K_D$ = 89 ± 13 nM, respectively. **C)** Binding isotherms of AP3-RhoBAST (linker) with SpyRho (10 nM, orange line) and GFP (5 nM, green line) giving dissociation constants of $K_D$ = 119 ± 15 nM and $K_D$ = 80 ± 16 nM respectively. Absorbance and fluorescence intensity measurements were performed in ASBT at 25°C. Minimal and maximal intensities were normalized to 0 and 1, respectively. $K_D$-values are given as mean ± s.d., $N$ = 3 individual experiments. **D, E, F)** Correlation of GFP and SpyRho fluorescence intensities in confocal images of HEK293T nuclei as shown in Fig. 1E ($N$ = 50 nuclei). HEK293T cells expressing H2B-GFP were fixed, permeabilized and visualized with AP3-RhoBAST:SpyRho using different fusion constructs (D) loop, (E) F30, (F) linker).

**G, H, I)** Colocalization test of GFP and SpyRho fluorescence in confocal images of HEK293T cells as shown in Fig. 1E using JACoP[19] to determine Pearson´s R values for different fusion constructs of AP3-RhoBAST (G) loop, (H) F30, (I) linker). **J)** Predicted secondary structure of AP3-RhoBAST (loop).

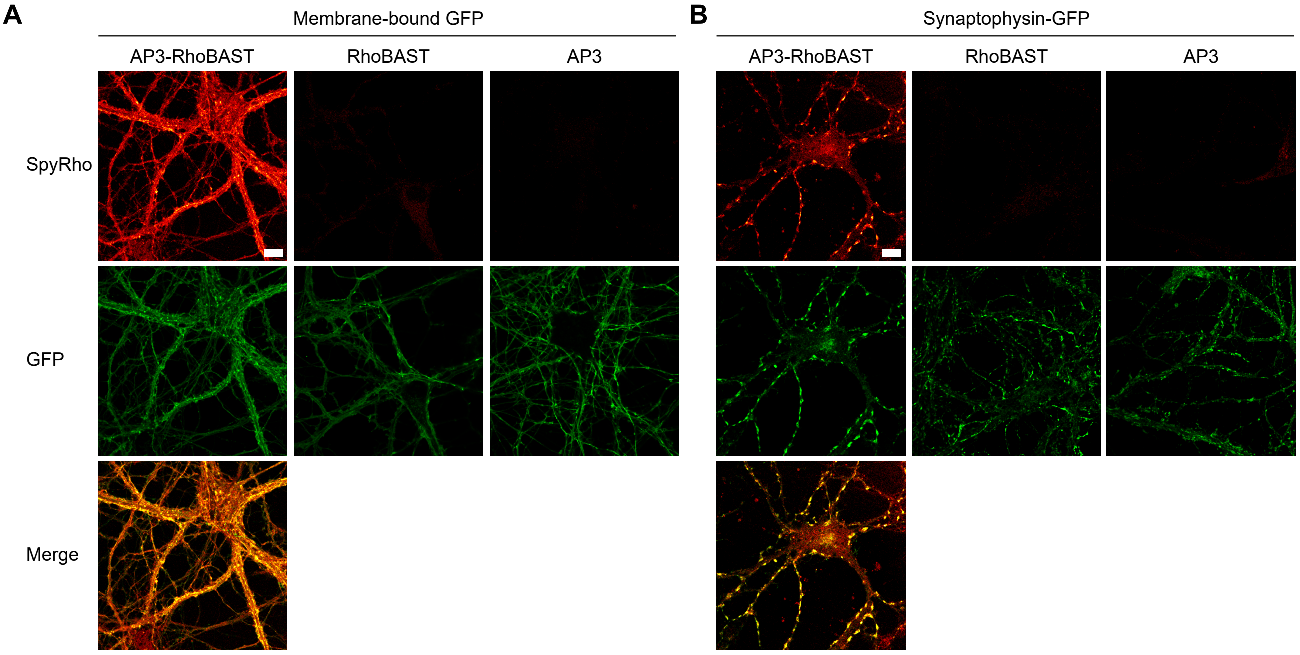


**Supplementary Figure S2: Confocal images of fixed hippocampal cells. Hippocampal cells expressing either A)** membrane-bound GFP tag or **B)** synaptophysin-GFP were fixed and permeabilized. Cells were incubated with 500 nM (bifunctional) aptamer, washed, and incubated with SpyRho (100 nM) prior to imaging. Successful staining was observed with AP3-RhoBAST, whereas RhoBAST and AP3, utilized as negative controls, exhibited no fluorescence signal. Scale bars, 10 µm.

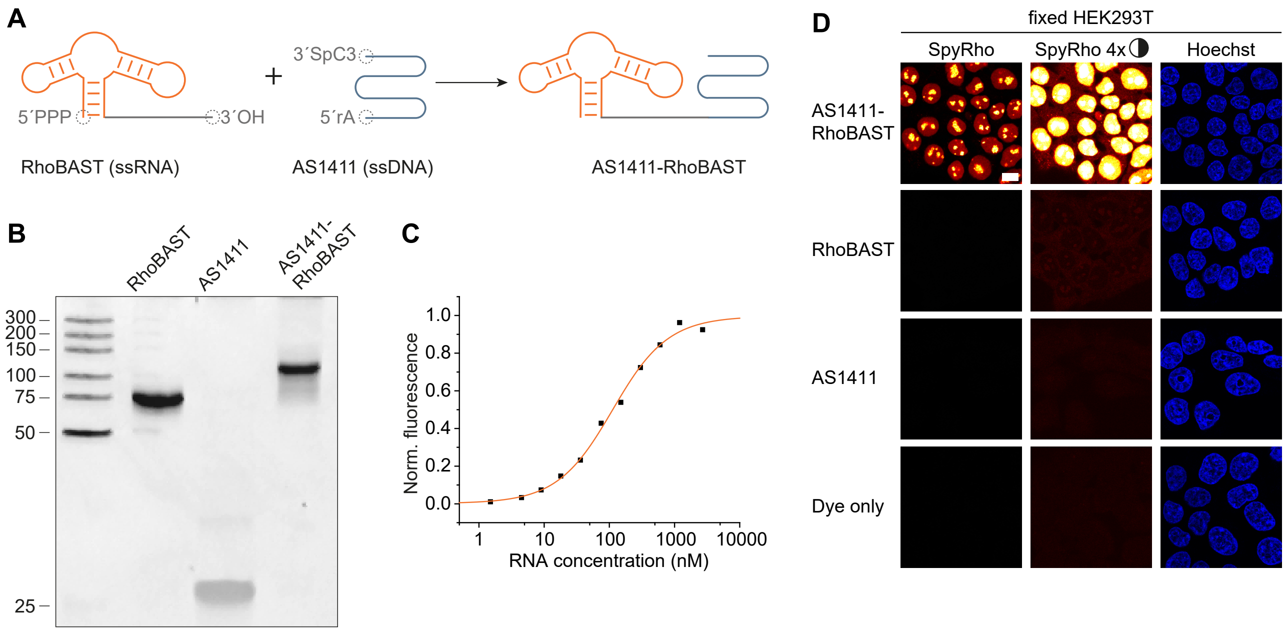


**Supplementary Figure S3: Synthesis, *in vitro* characterization, and application of AS1411-RhoBAST for confocal visualization of nucleolin in fixed HEK293T cells. A)** Schematic drawing of the ligation of *in vitro* transcribed RhoBAST with linker and single-stranded oligodeoxynucleotide AS1411 containing a 5´-adenosinmonophosphate (rA) and a 3´-spacer (3´-SpC3-OH) to form the DNA-RNA aptamer conjugate AS1411-RhoBAST. **B)** Analytical PAGE (20%) of RhoBAST with linker, AS1411, and the ligated DNA-RNA conjugate AS1411-RhoBAST stained with ethidium bromide. **C)** Binding isotherm of AS1411-RhoBAST with SpyRho (10 nM) giving a dissociation constant of $K_D$ = 113 ± 4 nM. Minimal and maximal fluorescence intensities were normalized to 0 and 1, respectively. $K_D$-values are given as mean ± s.d., $N$ = 3 independent experiments. **D)** Fixed HEK293T cells incubated with AS1411-RhoBAST to visualize nucleolin. As negative controls, RhoBAST, AS1411, and dye only are shown. The SpyRho channel is additionally shown with fourfold increased contrast settings. Scale bar, 10 µm.

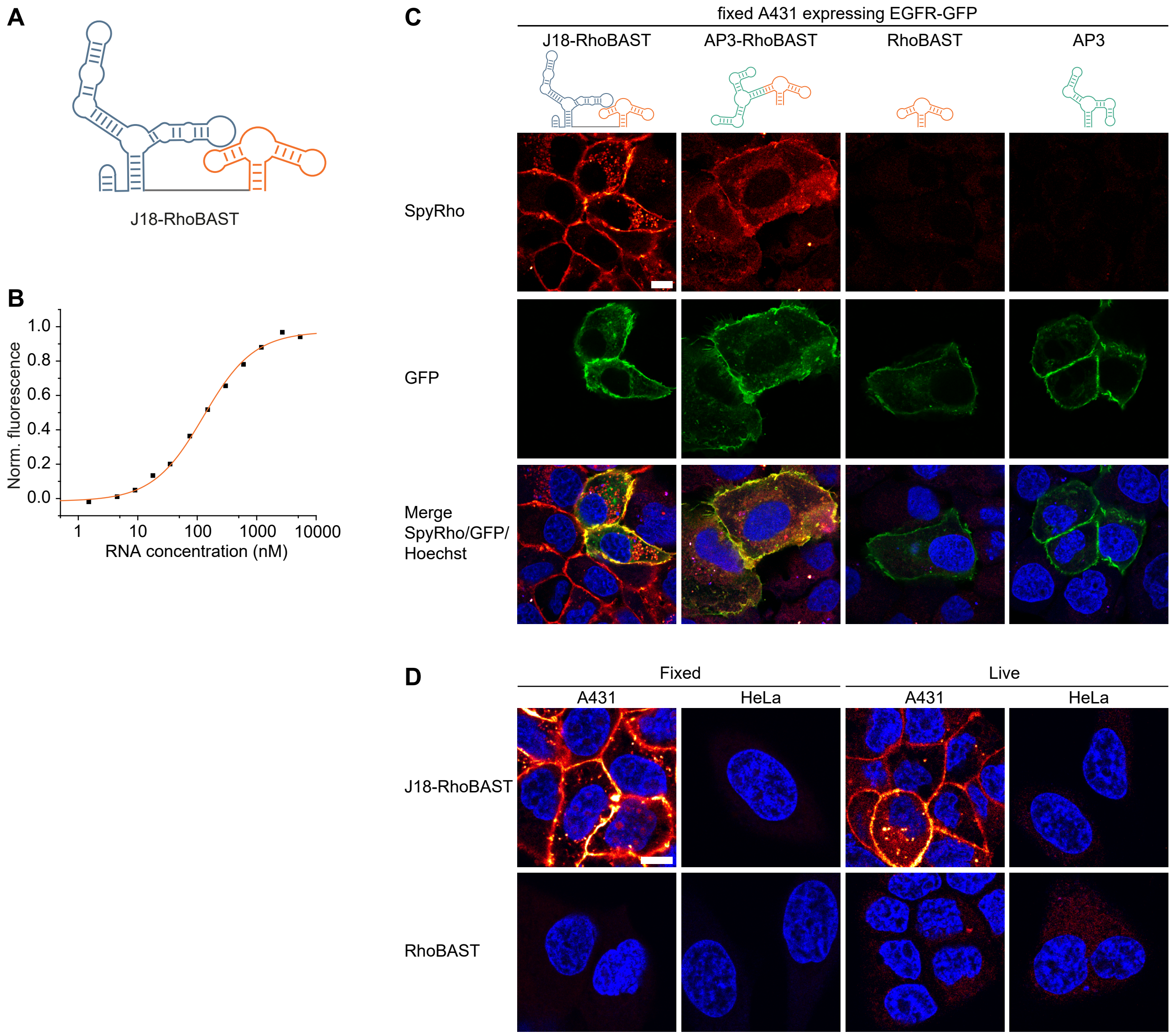


**Supplementary Figure S4: Characterization and application of J18-RhoBAST for confocal imaging of EGFR in fixed and on live cells. A)** Schematic drawing of bifunctional RNA aptamer J18-RhoBAST. **B)** Binding isotherm of J18-RhoBAST with SpyRho (10 nM) giving a dissociation constant of $K_D$ = 115 ± 9 nM. Fluorescence intensity measurements were performed in ASBT at 25°C. Minimal and maximal fluorescence intensities were normalized to 0 and 1, respectively. $K_D$-values are given as mean ± s.d., $N$ = 3 independent experiments. **C)** Transiently transfected A431 cells expressing EGFR-GFP were fixed, and EGFR was visualized either with J18-RhoBAST (500 nM, first column), staining all cells, or with AP3-RhoBAST (500 nM, second column), staining only the GFP-positive cells. As negative controls, the individual aptamers, RhoBAST (500 nM), and AP3 (500 nM), are shown. 100 nM SpyRho was applied. Scale bar, 10 µm. **D)** EGFR in fixed or on live A431 cells was visualized using J18-RhoBAST (1000 nM) with SpyRho (100 nM). As control fixed, and live HeLa cells were used. As negative control, RhoBAST (1000 nM, SpyRho 100 nM) is shown. Scale bar, 10 µm.

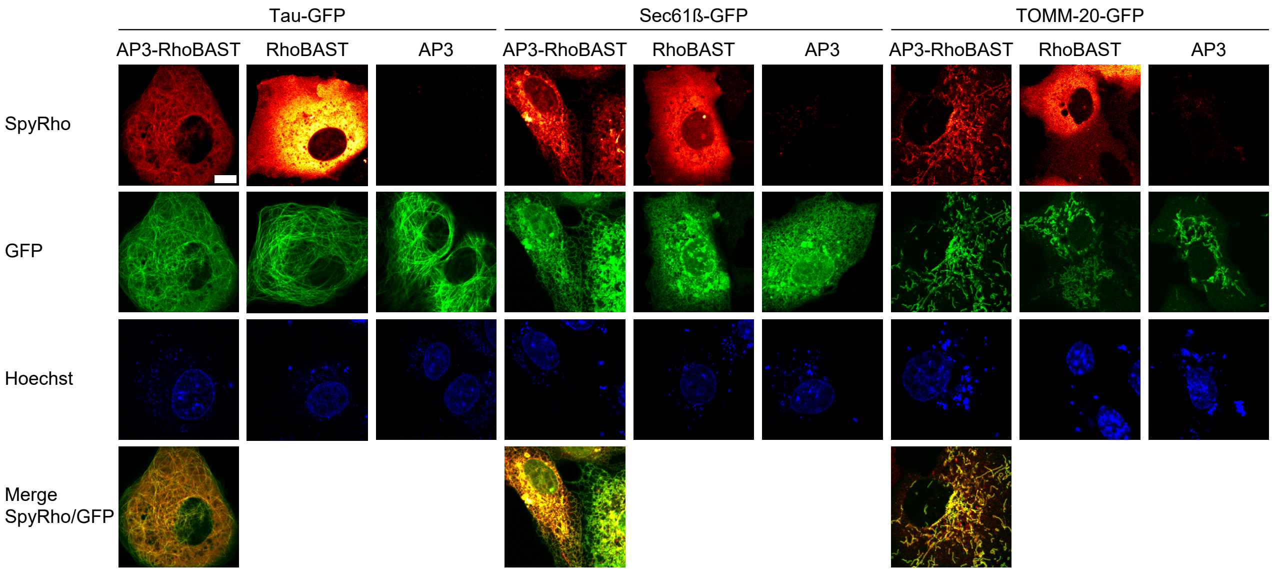


**Supplementary Figure S5: Confocal images of live COS7 cells co-expressing the genetically encoded aptamers in the Tornado expression system and the GFP-tagged target proteins.** COS7 cells were transiently co-transfected 48 h prior to imaging either with Tau-GFP, Sec61β-GFP or TOMM-20-GFP, and with Tornado-AP3-RhoBAST. As controls, Tornado-RhoBAST or Tornado-AP3 plasmids were used. The plasmids were applied in a 1:10 ratio (GFP-tagged POI to Tornado-aptamer). Live cells were incubated with 100 nM SpyRho for 30 minutes before imaging. Scale bar, 10 µm.

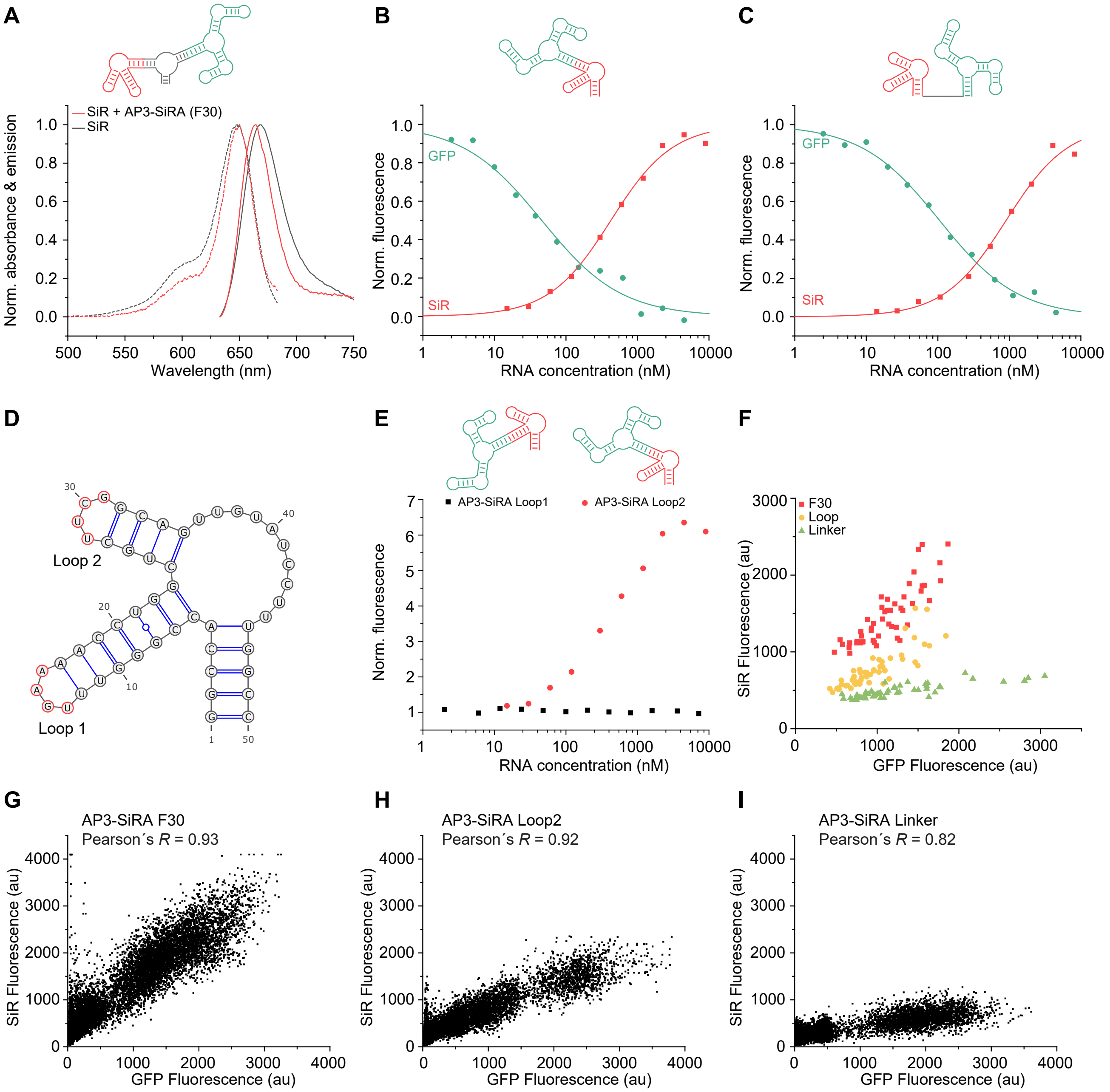


**Supplementary Figure S6: Comparison of different fusion constructs of bifunctional aptamer AP3-SiRA. A)** Normalized absorbance (dotted lines) and emission (solid line) spectra of SiR (200 nM) in the presence and absence of AP3-SiRA (5400 nM, F30). **B)** Binding isotherms of AP3-SiRA (loop2) with SiR (10 nM, red line) and GFP (5 nM, green line) yielding dissociation constants of $K_D$ = 360 ± 53 nM and $K_D$ = 63 ± 7 nM, respectively. **C)** Binding isotherm of AP3-SiRA (linker) with SiR (10 nM, red line) and GFP (5 nM, green line) yielding dissociation constants of $K_D$ = 1047 ± 123 nM and $K_D$ = 101 ± 12 nM, respectively. Absorbance and fluorescence intensity measurements were performed in ASBT at 25°C. Minimal and maximal fluorescence intensities were normalized to 0 and 1, respectively. $K_D$-values are given as mean ± s.d., $N$ = 3 independent experiments. **D)** Predicted secondary structure of SiRA, red regions indicate the tetraloops 1 and 2. **E)** Fluorescence intensity measurements of the titration of SiR (10 nM) with AP3-SiRA connected via the first or the second tetraloop. Intensity values were normalized to the fluorescence intensity of SiR in the absence of RNA. **F)** Correlation of GFP and SiR fluorescence intensities in confocal images of HEK293T nuclei as shown in Fig. 4D ($N$ = 50 nuclei). HEK293T cells expressing H2B-GFP were fixed, permeabilized and visualized with AP3-SiRA:SiR. **G, H, I)** Colocalization test of GFP and SiR fluorescence in confocal images of HEK293T cells as shown in Fig. 4D using JACoP[19] to determine Pearson´s R values for different fusion constructs of AP3-SiRA (G) loop, (H) F30, and (I) linker.

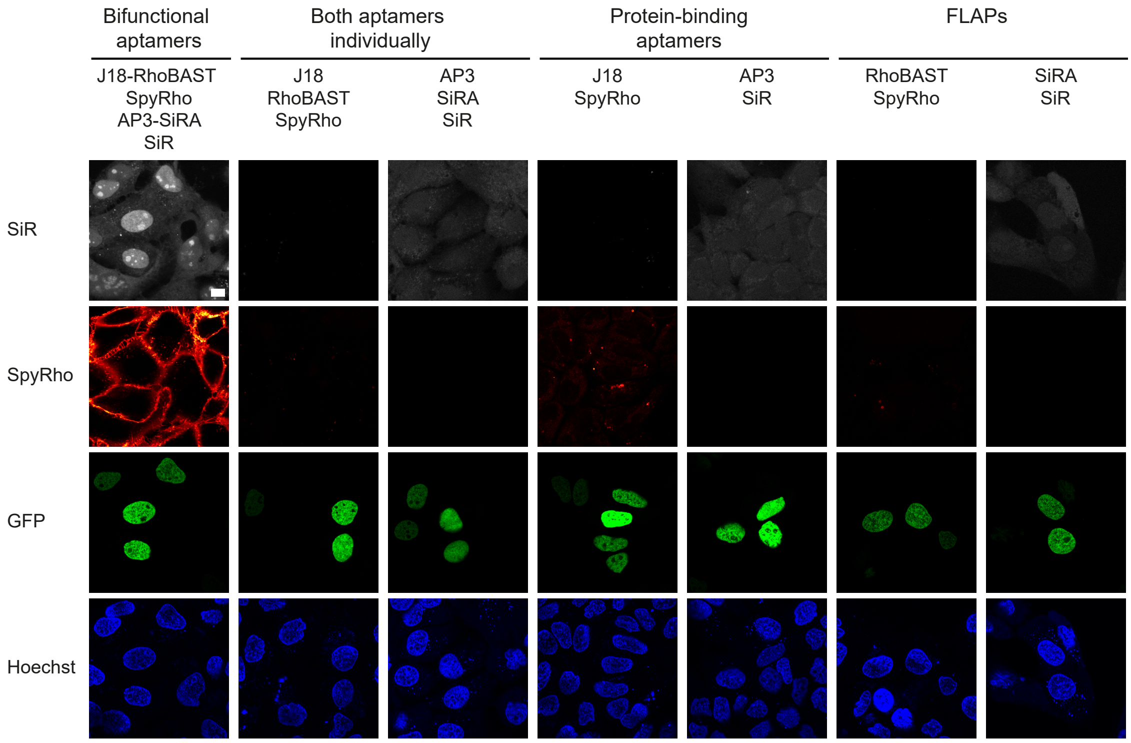


**Supplementary Figure S7: Dual-color confocal visualization of EGFR and H2B-GFP with J18-RhoBAST and AP3-SiRA, respectively.** Confocal images of A431 cells expressing H2B-GFP. First, EGFR was visualized in living A431 cells using J18-RhoBAST (500 nM) and SpyRho (100 nM). Then, the sample was fixed and H2B-GFP visualized using AP3-SiRA (1000 nM) and SiR (100 nM). As negative controls, the individual aptamers (J18, RhoBAST, AP3, SiRA), the protein-binding aptamers (J18, AP3), and the FLAPs (RhoBAST, SiRA) were applied with their cognate dye. Scale bar, 10 µm.

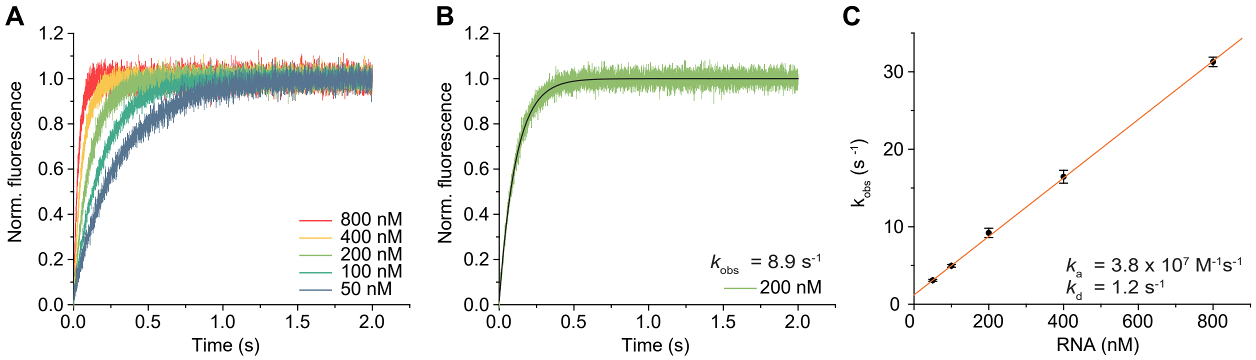


**Supplementary Figure S8: Determination of kinetic rate coefficients of AP3-RhoBAST using stopped-flow kinetic measurements. A)** Normalized fluorescence increase over time upon mixing AP3-RhoBAST (50, 100, 200, 400, 800 nM) and SpyRho (5 nM) in ASBT at 25°C. Data shown are averaged technical replicates (*N* = 5). **B)** Exponential fit of the fluorescence time curve from panel A (200 nM), giving the observed kinetic rate coefficient $k_{obs}$. **C)** Linear fitting of the observed kinetic rate coefficients $k_{obs}$ versus the RNA concentrations yields the association rate coefficient $k_a$ and dissociation rate coefficient $k_d$. Data points represent mean ± s.d. of three independent measurements.

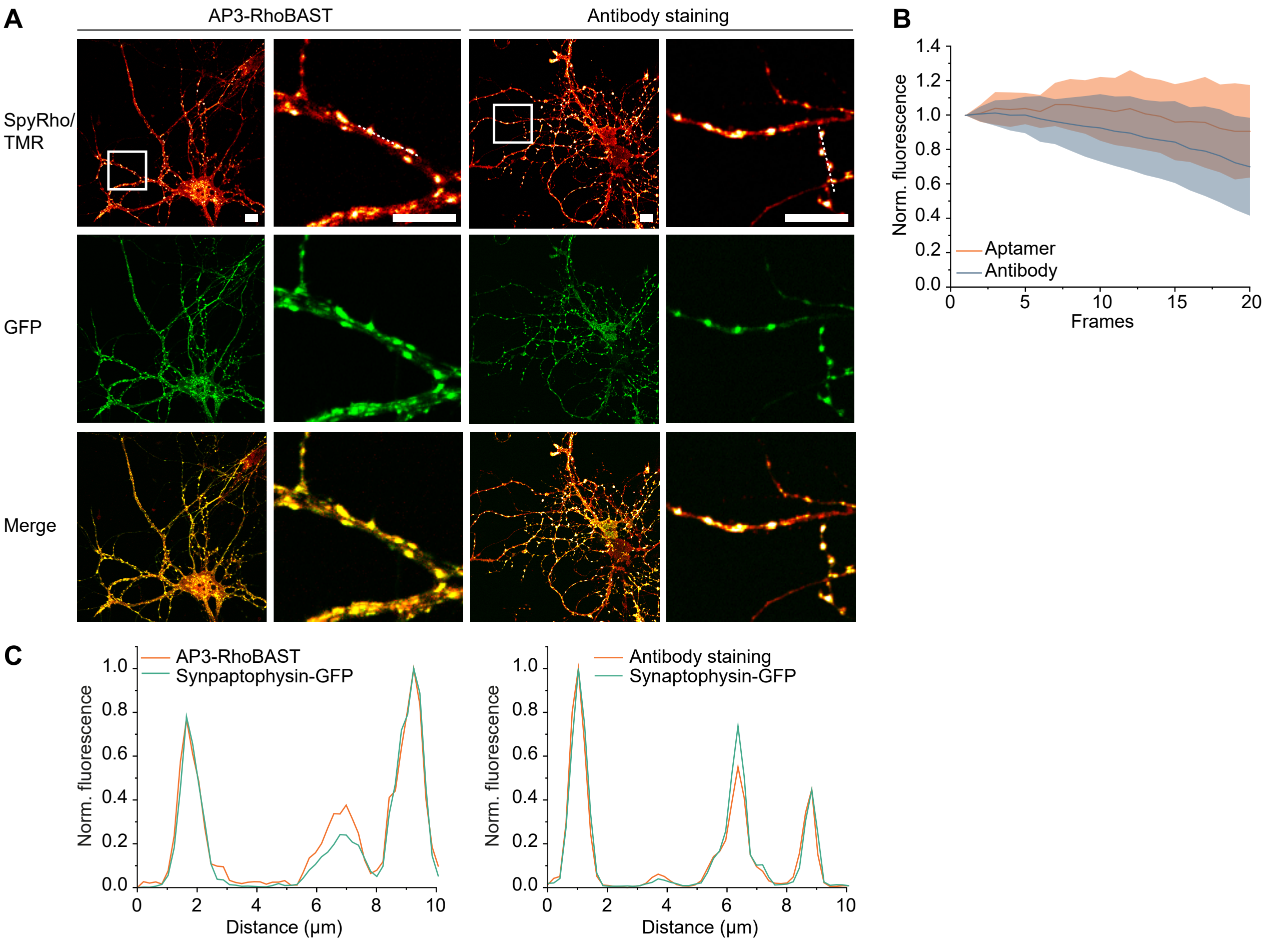


**Supplementary Figure S9: Comparison of synaptophysin visualization with bifunctional aptamer AP3-RhoBAST or with antibodies. A)** Hippocampal cells expressing GFP-tagged synaptophysin were fixed and permeabilized. Synaptophysin-GFP was either visualized with AP3-RhoBAST (500 nM) and SpyRho (100 nM) or antibodies using anti-synaptophysin primary antibody and tetramethylrhodamine-labelled secondary antibody. Note that images in the SpyRho/TMR channel are also shown in Figure 6. White frame indicates region of the zoom-in shown in the next column. Scale bars, 10 µm. **B)** Normalized fluorescence intensity of AP3-RhoBAST or antibody staining of GFP-tagged synaptophysin in cells shown in A) over 20 frames (mean ± s.d., $N$ = 30 synaptophysin clusters, laser power ~25 µW). Fluorescence intensity decreases by 9% in the bifunctional aptamer approach, whereas a 30% decrease was observed in antibody-stained cells. **C)** Normalized fluorescence profiles along the dashed lines in the close-ups shown in A).

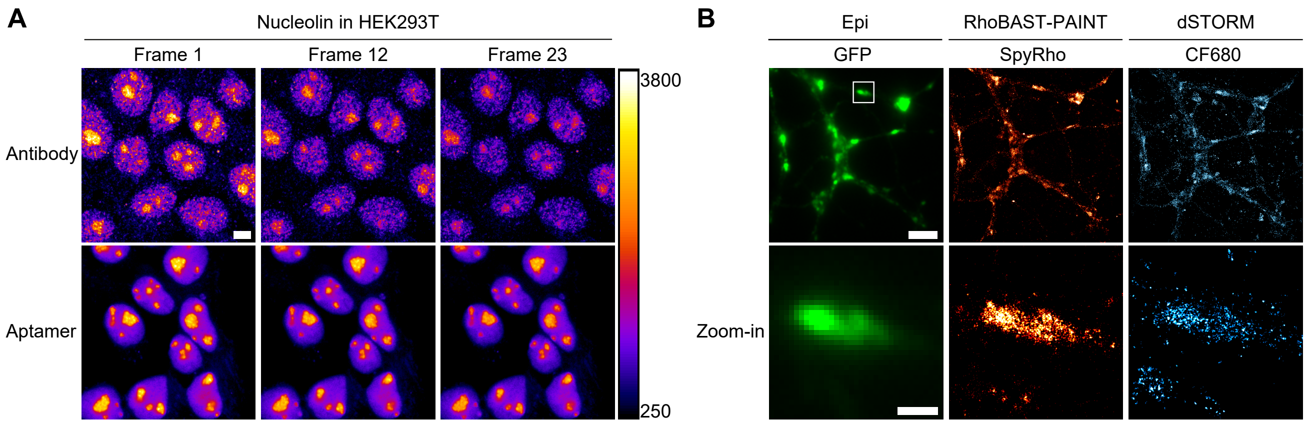


**Supplementary Figure S10: Comparison of bifunctional aptamers to antibodies in CLSM and SMLM. A)** Nucleolin in fixed and permeabilized HEK293T was visualized either with AS1411-RhoBAST:SpyRho or anti-nucleolin primary and rhodamine-labeled secondary antibody. Fluorescence decrease of aptamer or antibody staining of nucleolin shown for frame 1, 12, and 23 of confocal imaging using a laser power of 440 µW. Scale bar, 5 µm. The analysis of these images can be found in Figure 6C. **B)** Comparison of RhoBAST-PAINT and dSTORM in hippocampal cells expressing synaptophysin-GFP. Cells were fixed and permeabilized, incubated with AP3-RhoBAST (500 nM), washed with ASB, and imaged using SpyRho (1 nM). Reconstruction of 20,000 frames resulted in the super-resolved RhoBAST-PAINT image. Thereafter, the sample was washed with PBS and immunostained using anti-synaptophysin primary antibody and CF680-labeled secondary antibody. Under *d*STORM conditions, 20,000 frames were recorded and reconstructed. Scale bar, 5 µm. Scale bar of zoom-in, 1 µm.

**Supplementary Table S1:** *In vitro* properties of (bifunctional) aptamers. For SpyRho and SiR turn-on values are reported, whereas for GFP turn-off values are obtained. Values represent mean ± s.d. of three independent measurements.

| Aptamer | Fluorophore | Construct | $\lambda_{ex}$ [nm] | $\lambda_{em}$ [nm] | $K_D$ [nM] | Turn-on (off) [n-fold] |
|---|---|---|---|---|---|---|
| RhoBAST[2] | SpyRho | - | 562 | 581 | 34±2 | 60±18 |
| SiRA[1] | SiR | - | 648 | 668 | 430±70 | 7 |
| AP3[20] | GFP | - | 492 | 511 | 5.1 | 1.4 |
| AP3-RhoBAST | SpyRho | Linker | 562 | 581 | 119±15 | 50±4 |
| | | Loop | 562 | 581 | 107±17 | 47±2 |
| | | F30 | 562 | 582 | 122±10 | 41±4 |
| | GFP | Linker | 496 | 511 | 80±16 | 2.2±0.3 |
| | | Loop | 488 | 512 | 61±8 | 1.9±0.1 |
| | | F30 | 496 | 511 | 89±13 | 2.1±0.1 |
| AP3-SiRA | SiR | Linker | 649 | 663 | 1047±123 | 6.0±0.4 |
| | | Loop | 648 | 663 | 360±53 | 6.8±0.5 |
| | | F30 | 650 | 663 | 417±29 | 7.1±0.5 |
| | GFP | Linker | 497 | 511 | 101±12 | 1.4±0.1 |
| | | Loop | 495 | 510 | 63±7 | 1.5±0.1 |
| | | F30 | 496 | 511 | 105±18 | 1.4±0.1 |
| AS1411-RhoBAST | SpyRho | Linker | 562 | 582 | 113±4 | 37±3 |
| J18-RhoBAST | SpyRho | Linker | 562 | 581 | 115±9 | 21±1 |

**Supplementary Table S2:** Significance of quantification of AP3-FLAP performance shown in Figure 1D, E and Figure 4C, D. Two-sided two-sample *t*-test yielded the p-values. * p < 0.05, **** p < 0.001, n.s. not significant.

| Aptamer | Aptamer | p-value | Significance |
|---|---|---|---|
| AP3-RhoBAST (F30) | AP3-RhoBAST (Loop) | 0.04 | * |
| AP3-RhoBAST (F30) | AP3-RhoBAST (Linker) | 0.6 | n.s. |
| AP3-RhoBAST (Loop) | AP3-RhoBAST (Linker) | 0.05 | n.s. |
| RhoBAST | AP3-RhoBAST (F30) | $2.1 \times 10^{-37}$ | **** |
| RhoBAST | AP3-RhoBAST (Loop) | $6.3 \times 10^{-21}$ | **** |
| RhoBAST | AP3-RhoBAST (Linker) | $9.4 \times 10^{-34}$ | **** |
| AP3-SiRA (F30) | AP3-SiRA (Loop) | $1.1 \times 10^{-18}$ | **** |
| AP3-SiRA (F30) | AP3-SiRA (Linker) | $3.7 \times 10^{-33}$ | **** |
| AP3-SiRA (Loop) | AP3-SiRA (Linker) | $2.9 \times 10^{-11}$ | **** |
| SiRA | AP3-SiRA (F30) | $7.0 \times 10^{-43}$ | **** |
| SiRA | AP3-SiRA (Loop) | $2.5 \times 10^{-29}$ | **** |
| SiRA | AP3-SiRA (Linker) | $1.8 \times 10^{-41}$ | **** |

**Supplementary Table S3:** Kinetic rate coefficients for association ($k_a$) and dissociation ($k_d$) of aptamer:dye systems.

| **Aptamer:Dye** | **$k_a$ [$M^{-1}$ $s^{-1}$]** | **$k_d$ [$s^{-1}$]** |
|---|---|---|
| AP3-RhoBAST:SpyRho | $(3.77 \pm 0.04) \times 10^{7}$ | $1.21 \pm 0.06$ |
| RhoBAST:SpyRho[2] | $(2.1 \pm 0.1) \times 10^{7}$ | $1.8 \pm 0.1$ |
| AP3:GFP[20] | $5.9 \times 10^{5}$ | $2.66 \times 10^{-3}$ |

**Supplementary Table S4:** RNA and DNA (italic) sequences of used aptamer constructs.

| Construct | Sequence (5’ → 3’) |
|---|---|
| RhoBAST | GGAACCUCCGCGAAAGCGGUGAAGGAGAGGCGCAAGGUUAACCGCCUCAGGUUCC |
| AP3-RhoBAST (Linker) | GGAACCUCCGCGAAAGCGGUGAAGGAGAGGCGCAAGGUUAACCGCCUCAGGUUCCAAGAAAAGAAAGCUUCUGGACUGCGAUGGGAGCACGAAACGUCGUGGCGCAAUUGGGUGGGGAAAGUCCUUAAAAGAGGGCCACCACAGAAGCU |
| AP3-RhoBAST (Loop) | GGAACCUCCGCAGCUUCUGGACUGCGAUGGGAGCACGAAACGUCGUGGCGCAAUUGGGUGGGGAAAGUCCUUAAAAGAGGGCCACCACAGAAGCUGCGGUGAAGGAGAGGCGCAAGGUUAACCGCCUCAGGUUCC |
| AP3-RhoBAST (F30) | CUUGCCAUGUGUAUCGGUGGAACCUCCGCGAAAGCGGUGAAGGAGAGGCGCAAGGUUAACCGCCUCAGGUUCCUCCGAUACUCUGAUGAUGGGUAGCUUCUGGACUGCGAUGGGAGCACGAAACGUCGUGGCGCAAUUGGGUGGGGAAAGUCCUUAAAAGAGGGCCACCACAGAAGCUUCCCAUCAUUCAUGGCAAG |
| SiRA | GGCCACCGGGUUUGAAAACCUGGCUGCUUCGGCAGUUGUAUCCUUUGGCC |
| AP3-SiRA (Linker) | GGCCACCGGGUUUGAAAACCUGGCUGCUUCGGCAGUUGUAUCCUUUGGCCAAGAAAAGAAAGCUUCUGGACUGCGAUGGGAGCACGAAACGUCGUGGCGCAAUUGGGUGGGGAAAGUCCUUAAAAGAGGGCCACCACAGAAGCU |
| AP3-SiRA (Loop) | GGCCACCGGGUUUGAAAACCUGGCUGCAGCUUCUGGACUGCGAUGGGAGCACGAAACGUCGUGGCGCAAUUGGGUGGGGAAAGUCCUUAAAAGAGGGCCACCACAGAAGCUGCAGUUGUAUCCUUUGGCC |
| AP3-SiRA (F30) | CUUGCCAUGUGUAUCGGUGGCCACCGGGUUUGAAAACCUGGCUGCUUCGGCAGUUGUAUCCUUUGGCCUCCGAUACUCUGAUGAUGGGUAGCUUCUGGACUGCGAUGGGAGCACGAAACGUCGUGGCGCAAUUGGGUGGGGAAAGUCCUUAAAAGAGGGCCACCACAGAAGCUUCCCAUCAUUCAUGGCAAG |
| AP3 | AGCUUCUGGACUGCGAUGGGAGCACGAAACGUCGUGGCGCAAUUGGGUGGGGAAAGUCCUUAAAAGAGGGCCACCACAGAAGCU |
| J18-RhoBAST | GGCGCUCCGACCUUAGUCUCUGCAAGAUAAACCGUGCUAUUGACCACCCUCAACACACUUAUUUAAUGUAUUGAACGGACCUACGAACCGUGUAGCACAGCAGAAAAAAAGGGGAACCUCCGCGAAAGCGGUGAAGGAGAGGCGCAAGGUUAACCGCCUCAGGUUCCCC |
| J18 | GGCGCUCCGACCUUAGUCUCUGCAAGAUAAACCGUGCUAUUGACCACCCUCAACACACUUAUUUAAUGUAUUGAACGGACCUACGAACCGUGUAGCACAGCAGA |
| AS1411-RhoBAST | GGGGAACCUCCGCGAAAGCGGUGAAGGAGAGGCGCAAGGUUAACCGCCUCAGGUUCCCCAAGAAAAGAAA*GGTGGTGGTGGTTGTGGTGGTGGTGG* |
| AS1411 | *GGTGGTGGTGGTTGTGGTGGTGGTGG* |

**Supplementary Table S5:** DNA sequences of cloned pAV-U6+27-Tornado-Aptamer plasmids.

| Construct | Sequence (5' → 3') |
|---|---|
| pAV-U6+27-Tornado-RhoBAST | …GTCGACGGGCCGCACTCGCCGGTCCCAAGCCCGGATAAAATGGGAGGGGGCG<br>GGAAACCGCCTAACCATGCCGAGTGCGGCCGCAGGAACCTCCGCGAAAGCGGT<br>GAAGGAGAGGCGCAAGGTTAACCGCCTCAGGTTCCGTGGCCGCGGTCGGCGTG<br>GACTGTAGAACACTGCCAATGCCGGTCCCAAGCCCGGATAAAAGTGGAGGGTAC<br>AGTCCACGC… |
| pAV-U6+27-Tornado-AP3 | …GTCGACGGGCCGCACTCGCCGGTCCCAAGCCCGGATAAAATGGGAGGGGGCG<br>GGAAACCGCCTAACCATGCCGAGTGCGGCCGCAGCTTCTGGACTGCGATGGGAG<br>CACGAAACGTCGTGGCGCAATTGGGTGGGGAAAGTCCTTAAAAGAGGGCCACCAC<br>AGAAGCTCCGCGGTCGGCGTGGACTGTAGAACACTGCCAATGCCGGTCCCAAGC<br>CCGGATAAAAGTGGAGGGTACAGTCCACGC… |
| pAV-U6+27-Tornado-AP3-RhoBAST | …GTCGACGGGCCGCACTCGCCGGTCCCAAGCCCGGATAAAATGGGAGGGGGCG<br>GGAAACCGCCTAACCATGCCGAGTGCGGCCGCAGGAACCTCCGCAGCTTCTGGA<br>CTGCGATGGGAGCACGAAACGTCGTGGCGCAATTGGGTGGGGAAAGTCCTTAAA<br>AGAGGGCCACCACAGAAGCTGCGGTGAAGGAGAGGCGCAAGGTTAACCGCCTCA<br>GGTTCCGTGGCCGCGGTCGGCGTGGACTGTAGAACACTGCCAATGCCGGTCCCA<br>AGCCCGGATAAAAGTGGAGGGTACAGTCCACGC… |

**Supplementary Table S6:** Primary and secondary antibodies were used according to the supplier´s protocol in the listed dilution ratios.

| **Antibody** | **Dilution** | **Supplier** |
|---|---|---|
| Anti-EGFR (mouse, monoclonal, #MA5-13269) | 1:40 | Thermo Fisher Scientific |
| Anti-GFP (rabbit, monoclonal, recombinant, #G10362) | 1:100 | Thermo Fisher Scientific |
| Anti-H2B (rabbit, monoclonal, recombinant, #MA5-24697) | 1:1000 | Thermo Fisher Scientific |
| Anti-synaptophysin1 (rabbit, polyclonal, #101102) | 1:200 | Synaptic Systems |
| Anti-nucleolin (rabbit, polyclonal, #PA5-82860) | 1:50 | Thermo Fisher Scientific |
| Anti-rabbit IgG, Rhodamine (goat, #31670) | 1:200 | Thermo Fisher Scientific |
| Anti-mouse IgG (H+L), Rhodamine (goat, #31660) | 1:200 | Thermo Fisher Scientific |
| Anti-rabbit IgG (H+L), F(ab´) fragment, CF680 (goat, polyclonal, #SAB4600362) | 1:400 | Sigma-Aldrich |
